# Stress-Related Transcriptional Regulators Dominate the Conserved Core GRN for Three Cyanobacteria: Network Topology Maps the Highest-Influence Nodes as Promising Engineering Targets

**DOI:** 10.64898/2026.06.09.731130

**Authors:** Pavlo Bohutskyi, Rudolph DiMura, Zachary Johnson, Robert Li, David Anderson, Margaret Cheung

## Abstract

Cyanobacteria manage photosynthetic and environmental stresses through transcriptional programs controlled by regulators also affecting carbon flux, growth states, and metabolic output that bioproduction seeks to optimize. This regulatory architecture and its most influential nodes remain incompletely characterized. We hypothesized two influential regulator layers: a conserved core responding to common stresses, and species-specific regulators mediating strain-level niche adaptations. Mapping both layers underpins understanding genome→regulatory-network→phenotype flow, enabling global transcription machinery engineering for reliable bioproduction. To test our hypothesis, we constructed conserved-core and species-specific gene regulatory networks (GRNs) for three cyanobacteria, *Synechococcus elongatus* PCC 7942, *Synechocystis* sp. PCC 6803 and *Picosynechococcus* sp. PCC 7002, integrating a manually curated multi-pipeline regulator inventory with 1,098 harmonized transcriptome states for the 1,362-gene tri-homolog core genome. We quantified each regulator influence using local (degree, k-core), global (betweenness, closeness), and community-aware (eigenvector) centrality measures, and an Integrated Centrality score aggregating influence across complementary topological measures. High-influence regulators are predicted to exert broad metabolic effects when manipulated, making them priority candidates for single-target engineering interventions that modulate multiple genes and reprogram complex phenotypes. Across the three cyanobacteria, the two GRN layers proved topologically distinct: the conserved core concentrated influence in stress-related hubs (11 of its top 15 by Integrated Centrality were stress-related), while species-specific networks spread influence across functionally diverse regulators. Stress-coupled enrichment also held per individual centrality measure: regulators ranking top in both the core and species-specific GRNs by the same measure were mostly stress-related (15 of 19 instances), including the multi-stress regulators RpaB, Rre1, and BolA, the heat-shock HrcA, and the nitrogen NtcA. In species-specific GRNs, stress-related regulators remained the leading category alongside circadian, carbon-metabolism, morphology, and housekeeping regulators, including PlmA, Pex, TetR, and SrrB in PCC 7942; KaiC3, Sycrp1, Rre28, and Bhl in PCC 6803; and Zur, Sycrp1, and NarL in PCC 7002. High-influence putative regulators included the iron-stress AraC-family paralogs IutR1–IutR3, OmpR-family paralogs OmpR1–OmpR2, and chromosome- or plasmid-encoded Xre-family, AraC, and HypP. Stress regulation emerges as a recurring high-influence axis across these networks. The conserved core identifies universal regulatory programs, and species-specific layers reveal strain-level innovations for cross-strain transfer to support engineering of robust bioproduction.

**Importance:** Cyanobacteria are studied as platforms for sustainable, carbon-recycling production of fuels and chemicals from sunlight, water, and atmospheric carbon dioxide. Their reliable deployment in industrial settings is limited by environmental stresses that depress photosynthetic efficiency and product yields. The same regulatory proteins that govern stress responses also control how cells partition carbon, switch growth states, and direct metabolic output, making them natural levers for engineering robust production strains. Yet systematic, cross-species maps of these regulators have been missing. We present the first comparative regulatory map spanning three biotechnologically important model cyanobacteria, *Synechococcus elongatus* PCC 7942, *Synechocystis* sp. PCC 6803, and *Picosynechococcus* sp. PCC 7002, and identify the conserved regulators most influential across all three. The resulting catalog prioritizes candidate targets for experimental validation, and the supporting datasets and analytical framework are released for reuse to support efforts to engineer cyanobacterial strains for reliable industrial bioproduction.

## 1. Introduction

As primary producers in marine and freshwater ecosystems, cyanobacteria contribute a substantial fraction of global primary production (<u>1</u>), and their ability to convert atmospheric CO directly into biofuels and biochemicals using only sunlight and water positions them as platforms for sustainable, carbon-recycling bioproduction (<u>2</u>). Industrial-scale deployment requires strains that perform reliably under outdoor and industrial cultivation conditions, where high light, oxidative, osmotic, thermal, and nutrient stresses depress photosynthetic efficiency, destabilize cultures, and constrain yields. Stress regulation is therefore both a constraint and a tool for cyanobacterial engineering: the same regulators protecting cells under stress also govern carbon partitioning, growth-state transitions, and metabolic output, making them natural levers for engineering robust and productive strains. Yet our understanding of these regulators in cyanobacteria remains fragmented and largely strain-specific, with no systematic cross-species view of which regulators are most influential and how their stress-coupled functions are organized. Public RNA-seq repositories already hold the data to close that gap, and network-biology approaches now offer a viable comparative route.

Engineering such strains is challenging: stress tolerance and coordinated metabolic redirection for improved productivity may involve many genes across multiple pathways and feedback loops, and modifying single genes or single pathways is rarely sufficient. Global transcription machinery engineering (gTME) offers a complementary route: a single modification to a sigma factor or transcriptional regulator can reprogram broad swaths of the network and yield complex phenotypic changes (<u>3</u>, <u>4</u>). gTME has since produced diverse stress-tolerance phenotypes, including hyperosmotic, oxidative, and multi-stress robustness in *S. cerevisiae* (<u>5-7</u>), and has been extended to non-model bacterial and yeast hosts (<u>8-10</u>). In microalgae, transcription factor engineering has yielded strong bioproduction gains: overexpression of an endogenous bZIP regulator in *Nannochloropsis salina* raised fatty acid methyl ester content by 60% (<u>11</u>), and CRISPR knockout of a zinc-finger transcription factor in *Nannochloropsis gaditana* doubled lipid productivity by redirecting carbon from protein to lipid synthesis (<u>12</u>). Despite this progress in adjacent systems, gTME has not become part of the routine cyanobacterial metabolic engineering toolbox: existing work demonstrates single-regulator phenotypes rather than systematic, yield-oriented engineering campaigns, in part because the catalog of cyanobacterial transcriptional regulators is itself recent and still incomplete.

Two features of cyanobacterial transcription suggest this gap is closable. First, the core components of cyanobacterial transcription machinery are functionally cross-transferable: primary sigma factors function heterologously in phylogenetically close organisms (<u>13</u>), and cyanobacterial primary and alternative sigma factors recognize *Escherichia coli* consensus promoters and operate with *E. coli* RNA polymerase (<u>14</u>, <u>15</u>), as recently reviewed for cyanobacterial metabolic engineering (<u>16</u>). Second, the consensus binding motifs of upstream regulators including RpaA, RpaB, NtcA, NrrA, HrcA, and Rre1 are conserved across cyanobacteria and often shared with other bacteria, eukaryotic algae, and land plants (<u>17-19</u>). Together with existing engineering precedents in cyanobacteria, microalgae, and other microbes, this dual conservation makes cross-species transcription engineering biologically plausible. What has been missing is a systematic catalog of the most influential cyanobacterial regulators at two levels: those forming a conserved core shared across species and responding to common stresses, and those that are highly influential in specific species and enable niche-related functions due to evolutionary adaptations of individual strains.

Comparative computational analysis has become a productive route to bacterial biology, although progress has been uneven across scales. At the level of whole genomes, pan-genome and phylogenomic comparisons across bacterial taxa are now routine and have yielded robust evolutionary, taxonomic, and ecological insights in genera ranging from *Pseudomonas* to *Priestia*, and in phage populations associated with *Clostridioides difficile* (<u>20-22</u>). At the level of single bacterial species, machine-learning approaches applied to public RNA-seq compendia, including ICA-based iModulon decomposition and GENIE3-based GRN inference, have inferred transcriptional regulatory networks for *E. coli, B. subtilis, S. aureus, P. aeruginosa, P. syringae, P. putida, M. tuberculosis, S. enterica,* and *L. reuteri* (<u>23-25</u>); progress here is currently limited mainly by metadata, with fewer than 4% of microbial transcriptomic datasets in public repositories carrying annotations sufficient for high-throughput reanalysis (<u>26</u>). Progress is limited mainly by metadata, with <4% of public microbial transcriptomic datasets carrying annotations sufficient for high-throughput reanalysis (<u>26</u>). The closest prior efforts in cyanobacteria fall into three tiers: single-strain microarray compendia integrating heterogeneous datasets for PCC 6803, including a unified expression database and its meta-analysis revealing coordinated transcriptional adaptation across conditions (<u>27</u>, <u>28</u>); limited-scope cross-species expression datasets analyzed under specific stresses (<u>29</u>, <u>30</u>); and large RNA-seq compendia with genome-scale GRN reconstructions for individual strains, including iModulon- and GENIE3-based analyses of PCC 7942 circadian regulation (<u>31</u>, <u>32</u>), with reconstructions of Prochlorococcus MED4 (<u>33</u>) and an oxidative-stress-focused PCC 7942 network (<u>34</u>) currently underway. No integrated cross-species transcriptomic compendium or comparative GRN enabling regulator cataloging across *S. elongatus* PCC 7942, *Synechocystis* sp. PCC 6803, and *Picosynechococcus sp.* PCC 7002 has been published.

Identifying influential regulators requires cross-species comparison, which we develop here for three model cyanobacteria spanning much of the abiotic-stress and metabolic diversity relevant to biotechnological cultivation. *Synechococcus elongatus* PCC 7942 is a freshwater strain tolerant of high-light stress (<u>35</u>, <u>36</u>) and an obligate photoautotroph. *Synechocystis* sp. PCC 6803, a freshwater/brackish strain with substantial salinity tolerance (<u>37</u>), supports light-activated heterotrophic growth (<u>38</u>). *Picosynechococcus* sp. PCC 7002 is a marine strain notable for high-light tolerance, growth across a wide salinity range, and photomixotrophic capability (<u>39-43</u>). Phylogenetically, PCC 7002 and PCC 6803 are more closely related to one another than to PCC 7942 (<u>41</u>). To identify the most influential transcriptional regulators, we constructed core and species-specific gene regulatory networks (GRNs) from a curated multi-pipeline regulator inventory and a harmonized RNA-seq compendium, and quantified regulator influence using local (degree, k-core), global (betweenness, closeness), and community-aware (eigenvector) centrality measures. These measures together capture connectivity, information-flow position, and functional-module embedding rather than any single network feature; network-centrality analysis of this type extracts biologically meaningful regulatory architecture from cyanobacterial expression data despite the difficulty of predicting individual regulator–gene interactions (<u>32</u>). We define an Integrated Centrality score that aggregates these measures into a single ranking of regulators by joint topological influence.

Applying this comparative network framework to a compendium of nearly 1,100 expression states across the core genome (tripartite homologs) of PCC 7942, PCC 6803, and PCC 7002, we find that stress-response regulators are enriched among the high-influence positions of the conserved regulatory core. In contrast, in species-specific GRNs, stress-related regulators remain the leading category alongside carbon-metabolism, circadian, morphology, and housekeeping regulators. We then discuss how these findings can guide cyanobacterial transcription engineering across stress-tolerance, carbon-partitioning, and temporal-control applications. Together, this provides a systematic comparative analysis of cyanobacterial core regulatory architecture and a stress-anchored roadmap for engineering strategies that leverage stress-related regulators toward improved cyanobacterial bioproduction.

## 2. Methods

### 2.1. Assembling high-quality, species-specific RNA-seq compendia for *Synechococcus elongatus* PCC 7942, *Synechocystis* sp. PCC 6803, and *Picosynechococcus* sp. PCC 7002

We collected raw RNA sequencing data from the NCBI Sequence Read Archive (SRA) (<u>44</u>), Gene Expression Omnibus (GEO) (<u>45</u>), and the Joint Genome Institute databases (JGI) (<u>46</u>)) as of 31 January 2024. The overall initial dataset comprised 1,497 RNA-sequencing samples across 87 Bioprojects: 434 samples for PCC 7942, 776 samples for PCC 6803, and 287 samples for PCC 7002 (see **Supplementary Dataset S1**).

Raw RNA sequencing data alignment and gene quantification were performed using the *Rsubread* R package (<u>47</u>), mapping against the following genome assemblies: PCC 7942 – ASM1252v1; PCC 6803 – ASM972v1; and PCC 7002 – ASM1948v1 (<u>48</u>). We evaluated the resulting RNA-Seq datasets using *FastQC* (<u>49</u>), and manually refined them, selecting only data with well-described metadata and experimental conditions from the sourced databases or peer-reviewed publications.

To ensure high-quality final RNA-seq datasets, we implemented a rigorous three-step quality control and preliminary analysis process: i) we filtered out samples with insufficient sequencing depth (<100,000 protein-aligned reads); ii) Gene reads were normalized and transformed to log TPM, followed by global correlation evaluation among samples to detect and remove those with irregular expression patterns; iii) we excluded samples without biological replicates (except for time-series samples with adjacent neighbors showing high correlation) and samples with inadequate correlation coefficients (r ≤ 0.9) between replicates.

The resulting high-quality datasets, named Syn7942express, Syn6803express, and Syn7002express, contained 330, 548, and 220 RNA-Seq samples, respectively (see metadata and Log2 TPM gene counts for all sets in **Supplementary Datasets S2**).

### 2.2. Defining the conserved core genome and assembling the integrated cross-species RNA-seq compendium

To establish the core genome for PCC 7942, PCC 7002, and PCC 6803, we compared their genomes using the Proteome Comparison tool (<u>50</u>) available through the Bacterial and Viral Bioinformatics Resource Center (BV-BRC) (<u>51</u>). We identified conserved genes through protein sequence-based genome comparison using bidirectional BLASTP (<u>52</u>), with PCC 7942 as reference. Initial cutoffs were minimum coverage 30%, minimum identity 10%, and BLAST E-value <10 (**Supplementary Dataset S3-1**). The final tri-homologous and bi-homologous gene sets were defined by tightening these thresholds to minimum coverage 50% and minimum identity 40%, followed by manual curation (**Supplementary Dataset S3-2**).

We built a combined dataset by selecting orthologous genes from the species-specific Syn7942express, Syn6803express, and Syn7002express datasets and merging them into SynCOREexpress (**Supplementary Dataset S2**). This dataset contained expression for 1,362 orthologous genes under 1,098 unique expression states across diverse cultivation conditions, growth states, and environmental and genetic perturbations in PCC 7942, PCC 6803, and PCC 7002. We used this dataset for core gene regulatory network inference.

### 2.3. Identifying transcriptional regulators, assigning functional categories, and inferring core and species-specific Gene Regulatory Networks (GRNs)

To identify transcriptional regulators in PCC 7942, PCC 6803, and PCC 7002, we used four complementary computational pipelines: the Predicted Prokaryotic Transcription Factors (P2TF) database (<u>53</u>), the Encyclopedia of Well-Annotated DNA-binding Transcription Factors (ENTRAF) (<u>54</u>), the deep learning-based tool DeepTFactor (<u>55</u>), and the NCBI Prokaryotic Genome Annotation Pipeline (PGAP) (<u>56</u>). These pipelines integrate data from experimentally confirmed DNA-binding proteins in *E. coli* (RegulonDB; (<u>57</u>), *B. subtilis* (DBTBS; (<u>58</u>), and databases UniProt (<u>59</u>) and DNA-binding domain (DBD; (<u>60</u>) that predict regulators by sequence similarity through hidden Markov models (<u>61</u>) and convolutional neural networks (<u>55</u>).

We initially pooled all putative regulators predicted by the four pipelines (**Supplementary Dataset S4**), then conducted rigorous manual curation focused on regulators predicted by only one pipeline (**Supplementary Dataset S5**). Curation involved literature mining using PaperBLAST (<u>62</u>), functional domain identification via NCBI’s CDD (<u>63</u>), and protein similarity searches via NCBI’s BLAST (<u>64</u>). Regulators predicted by multiple pipelines were prioritized as more likely genuine regulators; single-method predictions received stringent examination (**Supplementary Dataset S5**).

To construct gene regulatory networks (GRNs), we used the random forest-based GENIE3 algorithm (<u>65</u>). GENIE3 quantified gene-to-gene expression associations, constrained by coupling expression matrices with the identified transcriptional regulators for each organism and the core genome. Output was a matrix of values representing regulatory interaction strengths between predicted regulators and all genes in the corresponding genomes. To focus on the most significant interactions, we pruned networks to ∼1,100 regulator-gene edges by removing the lowest-scoring. The lists of 889 genes (nodes), 38 transcription regulators (hub nodes), and 1,100 predicted TF–gene interactions (edges) are provided in **Supplementary Dataset S6**.

Functional category assignment to regulators. Each curated regulator was assigned to one of eight functional categories used throughout the figures and analyses: multi-stress response (light, redox, heat, osmotic), metal stress and homeostasis (Fe, Zn, Mn, heavy metals), nutrient stress and homeostasis (N, P), uncharacterized two-component stress response, circadian clock, carbon metabolism, cell morphology and motility, and housekeeping and transcription. Categories were defined and assigned by manual literature curation, supported by PaperBLAST homolog annotations (<u>62</u>) and conserved-domain analysis via NCBI’s CDD (<u>63</u>); **Tables 2** and **4** provide per-regulator functional descriptions and primary references that document the basis for each category assignment, including the stress-related functions. For regulators with documented multi-functional roles, the primary category was assigned based on the predominant function reported in the cyanobacterial literature or in the closest characterized homolog. For example, DnaA and NrdR were classified by their documented stress-coupled functions in cyanobacteria: DnaA couples replication initiation to heat-shock and oxidative stress (<u>66</u>, <u>67</u>), and NrdR couples ribonucleotide reductase regulation to DNA-damage and oxidative stress (<u>68</u>, <u>69</u>). Sigma factor names were unified across species using the SigA1, SigA4–SigA6 and SigF1, SigF2 convention to enable direct cross-species comparison in **Figures 3** and **5**; species-specific literature naming (RpoD1–RpoD4 in PCC 7942) is provided in the figure captions.

### 2.4. Detecting Louvain communities in GRNs and identifying enriched COG functional categories and KEGG pathways

We utilized *Networkx* (<u>70</u>) to perform Louvain clustering, identifying gene communities associated with specific regulators. To assess overlap between gene sets of different regulators, we performed hierarchical clustering on Jaccard distance: a pairwise Jaccard matrix across regulator gene-target sets followed by average-linkage clustering. Functional enrichment used Fisher’s exact test with Benjamini-Hochberg FDR adjustment for Q-value computation. We used the Clusters of Orthologous Groups (COG; (<u>71</u>) and Kyoto Encyclopedia of Genes and Genomes (KEGG; (<u>72</u>) databases to define functional enrichments for Louvain clusters and predicted regulons. KEGG pathways were visualized with Pathview (<u>73</u>). Enrichment p-values were calculated using the Fisher’s exact test hypergeometric formula (Eq. 1):

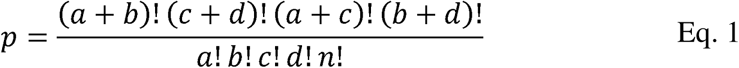

where: **a** corresponds to the number of genes in a cluster which overlap with the functional gene set of interest, **b** corresponds to the number of genes in the cluster which are not in the predicted gene sets. **c** corresponds to the number of genes in the background gene set not in the predicted cluster, and **d** corresponds to the background genes, all genes not in the predicted regulon or a gene set. COG enrichments were considered significant at a *q-value* threshold of 0.05, while KEGG pathway enrichments were evaluated at a q-value threshold of 0.01.

### 2.5. Identifying highest-influence regulators by centrality analysis and cross-validating across conserved and species-specific GRNs

To identify the highest-influence regulators, we conducted topological analysis of the GRNs. We used NetworkX (<u>70</u>) to determine GRN properties including eigenvector centrality and k-core. Network visualization and additional centrality measures (betweenness, closeness, degree, stress) were calculated using Cytoscape (<u>74</u>).

To integrate these measures into a unified regulator-influence ranking, we developed the Integrated Centrality (IC) score (Eq. 2), combining six individual network topology measures each normalized to its highest value. This approach assesses each regulator’s relative importance across multiple topology and centrality dimensions.

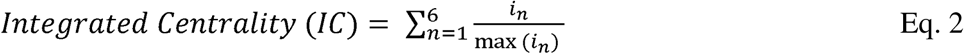

where *i* represents one of six centrality measures (k-core, betweenness, closeness, degree, stress, eigenvector) for a given regulator node.

#### Top-regulator selection and cross-validation across GRNs

For each individual centrality measure (degree, k-core, betweenness, closeness, stress, eigenvector), the top regulators of each species-specific GRN were defined as those ranking in the top decile by that measure, capped at 10 regulators per measure. For the conserved core GRN, top regulators were defined more stringently as the top three within the top decile by each measure, to focus the cross-species analysis on the most consistently influential nodes. Integrated Centrality top-regulator sets aggregate regulators identified across the six individual measures. To assess conservation of high-influence regulators across species, we cross-validated top regulators of the core GRN against top regulators of each species-specific GRN by the same measure: a regulator was counted as cross-validated for a given (species, measure) pair when its ortholog (defined by the bidirectional BLASTP homology above) ranked in the top three of the core GRN and also in the top decile of the species-specific GRN by that same measure. Cross-validated regulators are reported by (species × measure) cell in **Figure 4** (19 instances representing 15 distinct regulators). Species-specific top regulators (**Figure 5**) were defined as regulators in the top decile of a species-specific GRN by a given measure whose orthologs did not rank in the top three of the core GRN by the same measure (111 cell entries representing 39 distinct regulators). In **Figures 4** and **5**, regulator symbols are sized proportionally to the normalized centrality score within each cell and ordered top-to-bottom by descending score.

## 3. Results and Discussion

We conducted a comparative genomic analysis of three biotechnologically important cyanobacteria with well-characterized genomes and diverse physiology (<u>43</u>, <u>75</u>, <u>76</u>): *Synechococcus elongatus* PCC 7942 (PCC 7942), *Synechocystis* sp. PCC 6803 (PCC 6803), and *Picosynechococcus* sp. PCC 7002 (PCC 7002). These strains inhabit distinct ecological niches: PCC 7942 freshwater (<u>35</u>), PCC 7002 marine (<u>39</u>), and PCC 6803 likely brackish origin (<u>37</u>), yet all three tolerate high light conditions, a trait especially pronounced in PCC 7002 (<u>36</u>, <u>40</u>, <u>43</u>). To examine conserved and species-specific elements of their transcriptional regulatory architecture, we first defined the shared core genome, then assembled cross-species expression resources, predicted transcriptional regulators, inferred core and species-specific GRNs, and applied network topology analysis to identify shared and strain-specific high-impact regulators.

### 3.1. Composition and functional analysis of the conserved core genome across three model cyanobacteria

Using PCC 7942 as the reference, we identified homologous genes in the other two strains by protein sequence identity and coverage thresholds, revealing a conserved core genome of 1,362 genes shared across all three strains (**Figure 1A**, complete list in Supplementary **Dataset S3**). This tri-homologous core represents ∼50% of total genes in PCC 7942 and PCC 7002 but only ∼33% in PCC 6803, whose larger genome accommodates a substantially expanded strain-specific gene set. Pairwise duo-homolog sets reflect the underlying phylogeny: 483 genes are shared between PCC 6803 and PCC 7002, 160 between PCC 7942 and PCC 6803, and 140 between PCC 7942 and PCC 7002, consistent with prior reports that PCC 7002 and PCC 6803 are more closely related to each other than to PCC 7942 (<u>41</u>), and with their shared ability to tolerate high salinity (<u>77</u>) and utilize organic carbon (<u>38</u>, <u>42</u>). PCC 6803 supports light-activated heterotrophic growth (<u>38</u>) and PCC 7002 supports photomixotrophic growth (<u>42</u>), whereas PCC 7942 is restricted to photoautotrophy. The largest duo-homolog set (PCC 6803–PCC 7002) showed no significant functional enrichment, but the two smaller sets carried distinct functional signatures: the PCC 7942–PCC 7002 duo-homologs were enriched for secondary metabolite biosynthesis, transport, and catabolism (COG category Q), cysteine and methionine biosynthesis (KEGG ko00270), and quorum sensing (KEGG ko02024), while the PCC 7942–PCC 6803 duo-homologs were enriched for porphyrin metabolism (KEGG ko00860) and streptomycin biosynthesis (KEGG ko00521).

**Figure 1.**
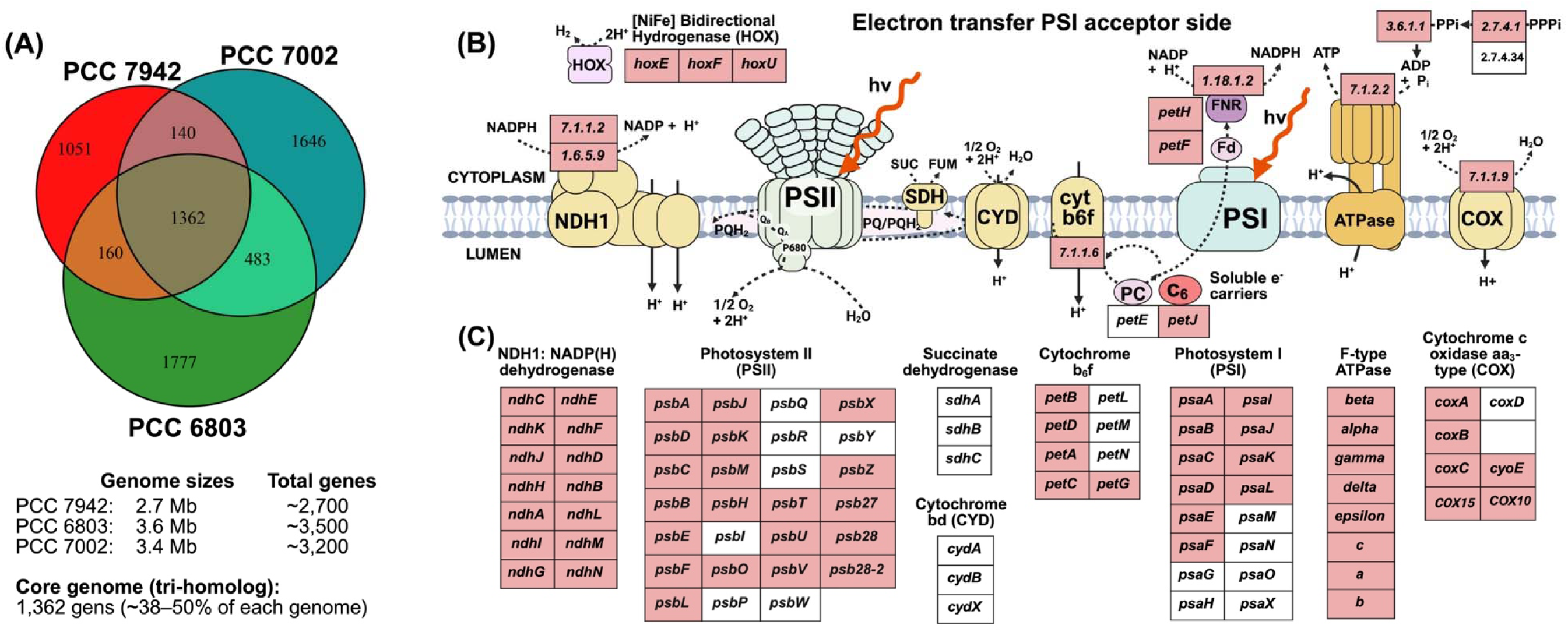
Genome-wide and energy-metabolism gene conservation across three model cyanobacteria. **(A)** Three-strain Venn diagram of homologous gene sets across *S. elongatus* PCC 7942, *Synechocystis* sp. PCC 6803, and *Picosynechococcus* sp. PCC 7002. Numbers indicate tri-homologs at the central intersection (1,362 genes), pairwise duo-homologs, and strain-specific singletons; genome sizes are listed below. **(B)** Schematic of the cyanobacterial thylakoid membrane integrating photosynthetic and respiratory electron transport, including the [NiFe] bidirectional hydrogenase (HOX), NADP(H) dehydrogenase (NDH-1), photosystem II (PSII), succinate dehydrogenase (SDH), cytochrome bd quinol oxidase (CYD), cytochrome b6f, photosystem I (PSI), F-type ATPase, and aa3-type cytochrome c oxidase (COX). Soluble electron carriers are shown for the cytoplasmic acceptor side of PSI (ferredoxin, Fd; ferredoxin-NADP+ reductase, FNR) and the lumenal donor side of PSI (plastocyanin, PC; cytochrome c6, C6). EC numbers (italicized) are shown for major catalytic activities. **(C)** Gene-level conservation in each complex from panel B. Pink boxes denote tri-homologous genes conserved across all three strains; white boxes denote genes non-homologous in at least one strain. Photosynthetic complexes (PSII, PSI, cytochrome b6f, F-type ATPase) are largely tri-homologous, with strain-specific exceptions concentrated in accessory subunits (*psbI*, *psbP*, *psbQ*, *psbR*, *psbS*, *psbW*, *psbY* of PSII; *petL*, *petM*, *petN* of cyt b6f; *psaG*, *psaH*, *psaM*, *psaN*, *psaO*, *psaX* of PSI; *coxD* of cytochrome c oxidase). Respiratory complexes show greater partial conservation, with succinate dehydrogenase (*sdhA*, *sdhB*, *sdhC*) and cytochrome bd quinol oxidase (*cydA*, *cydB*, *cydX*) absent or non-homologous in at least one strain.

Functional enrichment analysis of the 1,362-gene tri-homologous core revealed strong conservation of essential cellular machinery (**Table 1**). COG analysis identified significant enrichment in Translation (category J) and Energy production and conversion (C), and KEGG pathway analysis identified Biosynthesis of secondary metabolites (ko01110) and Ribosome (ko03010) as the most enriched categories. The large number of homologous ribosomal proteins was expected, given their high sequence conservation across bacterial species and even domains despite considerable structural diversity (<u>78</u>, <u>79</u>). Both analyses highlighted strong conservation of energy-metabolism genes operating under light (photosynthesis) and dark conditions (oxidative phosphorylation), examined below. Beyond translation and energy metabolism, the core was enriched for amino-acid metabolism (COG category E; KEGG ko00250) and nucleotide metabolism and transport (COG category F), with 19 homologous genes in glycine-serine-threonine metabolism and 15 each in alanine-aspartate-glutamate and cysteine-methionine metabolism.

**Table 1.** Functional enrichment of tri-homologous and duo-homologous gene sets across *Synechococcus elongatus* PCC 7942, *Synechocystis* sp. PCC 6803, and *Picosynechococcus* sp. PCC 7002. Enriched Clusters of Orthologous Groups (COG) categories and Kyoto Encyclopedia of Genes and Genomes (KEGG) pathways are listed for the 1,362-gene tri-homologous core (genes shared by *Synechococcus elongatus* PCC 7942, *Synechocystis* sp. PCC 6803, and *Picosynechococcus* sp. PCC 7002) and for the two duo-homolog sets that showed significant enrichment (PCC 7942–PCC 7002 and PCC <u>7942–PCC 6803).</u>

| Homology | COG | KEGG <sup>A</sup> | Description | Gene ratio <sup>B</sup> | *p*-value |
| --- | --- | --- | --- | --- | --- |
| PCC 7942 – | J | – | Translation | 148/1362 | 5.17e-21 |
| PCC 6803 – | C | – | Energy Production and Conversion | 144/1362 | 1.60e-5 |
| PCC 7002 | F | – | Nucleotide Metabolism and Transport | 75/1362 | 2.64e-5 |
|  | H | – | Coenzyme Metabolism | 108/1362 | 5.58E-04 |
|  | O | – | Post-translational modification, protein turnover, chaperone functions | 64/1362 | 3.22E-03 |
|  | E | – | Amino Acid metabolism and transport | 84/1362 | 6.34E-03 |
|  | I | – | Lipid Metabolism | 36/1362 | 1.53E-02 |
|  | – | ko01110 | Biosynthesis of Secondary Metabolites | 203/1362 | 7.07e-27 |
|  | – | ko03010 | Ribosome | 53/1362 | 8.23e-17 |
|  | – | ko01230 | Biosynthesis of Amino Acids | 81/1362 | 1.00e-13 |
|  | – | ko00195 | Photosynthesis | 49/1362 | 1.20e-7 |
|  | – | ko00970 | Aminoacyl-tRNA Biosynthesis | 25/1362 | 4.00e-7 |
|  | – | ko01120 | Microbial Metabolism in Diverse Environments | 87/1362 | 1.03e-6 |
|  | – | ko00190 | Oxidative Phosphorylation | 40/1362 | 4.42e-6 |
|  | – | ko00230 | Purine Metabolism | 45/1362 | 6.16e-6 |
|  | – | ko01200 | Carbon Metabolism | 53/1362 | 8.76e-6 |
| PCC 7942 –<br>PCC 7002 | Q | – | Secondary Metabolite Biosynthesis, Transport, and Catabolism | 8/140 | 1.75e-3 |
|  | – | ko02024 | Quorum Sensing | 8/140 | 4.45e-4 |
|  | – | ko00270 | Cysteine and Methionine Metabolism | 7/140 | 1.16e-3 |
| PCC 7942 – | – | ko00521 | Streptomycin Biosynthesis | 6/160 | 4.66e-5 |
| PCC 6803 | – | ko00860 | Porphyrin Metabolism | 10/160 | 3.57e-4 |

**Gene ratio** <sup>(B)</sup>: number of genes in the enriched category divided by the total genes in the homolog set (e.g., 148/1362 for COG category J in the tri-homologous core).
**KEGG pathways** <sup>(A)</sup>: the top ten KEGG categories ranked by *p*-value are listed for each gene set.
The PCC 6803–PCC 7002 duo-homolog set (483 genes), the largest pairwise homolog group, showed no statistically significant functional enrichment and is not listed.

Energy-metabolism genes encoding the photosynthetic and respiratory electron transport machinery were strongly tri-homologous, with strain-specific exceptions concentrated in accessory subunits and alternative oxidases (**Figure 1B, 1C**). The core photosynthetic complexes (photosystem II/PSII, photosystem I/PSI, cytochrome b6f, F-type ATPase) and the photosynthetic electron carriers ferredoxin (Fd), ferredoxin-NADP+ reductase (FNR), and cytochrome c6 (encoded by *petJ*) were conserved across all three strains, as were genes for porphyrin metabolism and carotenoid biosynthesis, indicating maintained alternative light-harvesting capacity and photoprotection (<u>80</u>). Strain-specific differences in photosynthesis localized to accessory subunits: *psbI* (PSII reaction center protein I) and *psbP* (oxygen-evolving complex 23K protein) were shared between PCC 6803 and PCC 7002 but absent in PCC 7942, as was *psaM* (PSI reaction center subunit XII), while the cytochrome b6f subunit *petL* and the plastocyanin gene *petE* were shared between PCC 6803 and PCC 7942 but absent in PCC 7002. Respiratory machinery showed a similar tri-homology-with-exceptions pattern. NADP(H) dehydrogenase (NDH-1), aa3-type cytochrome c oxidase, the heme A biogenesis genes COX15 and COX10 (*ctaA* and *ctaB*), and the F-type ATPase were tri-homologous across the three strains, while three respiratory complexes lacked tri-homology: succinate dehydrogenase (*sdhA*, *sdhB*, *sdhC*) was shared between PCC 6803 and PCC 7002 but absent in PCC 7942; cytochrome bd quinol oxidase (*cydA*, *cydB*) was shared between PCC 6803 and PCC 7942 but absent in PCC 7002; and cbb3-type cytochrome c oxidase was unique to PCC 7942. These strain-specific exceptions may reflect adaptations to different environmental light regimes and energy requirements.

Together, these findings define a 1,362-gene tri-homologous core that is enriched for translation, energy metabolism, and amino-acid metabolism, alongside duo-homolog and strain-specific gene sets reflecting the distinct phylogeny and metabolic capabilities of each strain. With the conserved core genome defined, we next turned to the transcriptional machinery that regulates these shared genes.

### 3.2. Multi-pipeline computational prediction and manual curation of conserved and species-specific transcriptional regulators across three model cyanobacteria

DNA-binding transcription factors (TFs) and sigma factors are the major elements of bacterial transcription. In model cyanobacteria, while individual regulators have been extensively studied, regulator inventories remain incomplete and the relative network influence of individual regulators across conserved and species-specific layers has not been systematically resolved. For comprehensive regulator coverage across PCC 7942, PCC 7002, and PCC 6803, we applied four complementary computational pipelines: the deep learning-based tool DeepTFactor (<u>55</u>), the Predicted Prokaryotic Transcription Factors (P2TF) database (<u>53</u>), the Encyclopedia of Well-Annotated DNA-binding Transcription Factors (ENTRAF, (<u>54</u>)), and functional annotation from NCBI’s Prokaryotic Genome Annotation Pipeline (<u>56</u>, <u>81</u>).

The initial computational predictions yielded large putative TF sets across the three strains (140, 176, and 216 candidates for PCC 7942, PCC 7002, and PCC 6803, respectively; **Figure 2A** and **2B**, complete list in **Supplementary Dataset S4**), but with substantial discrepancies between pipelines: many candidates were predicted by only a single method. These pipeline-level disagreements motivated a rigorous manual curation of the combined initial set.

**Figure 2.**
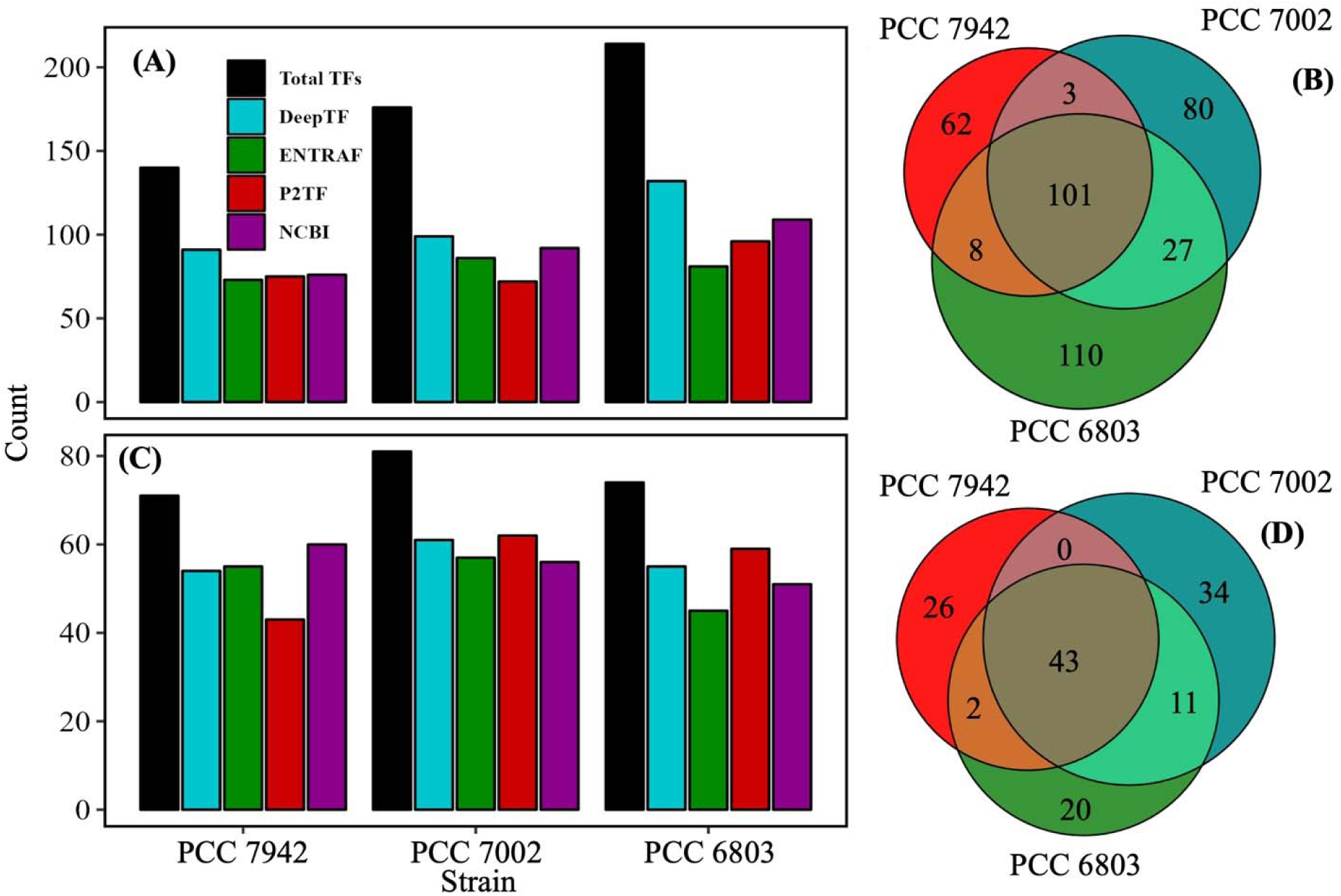
Multi-pipeline computational prediction and manual curation of transcriptional regulators across *S. elongatus* PCC 7942, Synechocystis sp. PCC 6803, and Picosynechococcus sp. PCC 7002. (A) Initial putative regulator counts predicted by four computational pipelines (DeepTFactor, ENTRAF, P2TF, NCBI PGAP) for each strain; black bars indicate combined totals (140, 177, and 216 for PCC 7942, PCC 7002, and PCC 6803). (B) Cross-strain overlap of initial putative regulators (101 shared across all three strains). (C) Final curated regulator counts per pipeline (71, 81, and 74 for PCC 7942, PCC 7002, and PCC 6803). (D) Cross-strain overlap of final curated regulators, with 43 conserved across all three strains as orthologs.

Manual curation focused on single-pipeline candidates, applying literature mining via PaperBLAST (<u>62</u>) and conserved-domain analysis via CDD (<u>82</u>) to evaluate functional annotations of each candidate and close homologs. Candidates supported by multiple pipelines were retained as high-confidence regulators, while single-pipeline candidates were retained only when literature and domain evidence confirmed a transcriptional regulatory function. Curation discarded two false-positive classes consistently across all three genomes: (i) candidates lacking identifiable DNA-binding domains (response regulators without cognate DNA-binding output, toxin-antitoxin components, hypothetical proteins, and proteins with domains of unknown function), and (ii) candidates with DNA-binding domains that function in DNA replication, repair, or transposition rather than transcription. The latter class was particularly notable in PCC 6803, whose genome carries a substantially expanded set of transposase genes that DeepTFactor frequently flagged as putative TFs based on their DNA-binding domains alone.

After curation, the final regulator sets comprised 71, 81, and 74 transcriptional regulators for PCC 7942, PCC 7002, and PCC 6803, respectively (**Figure 2C** and **2D**, complete list in **Supplementary Dataset S5**), encompassing DNA-binding TFs, sigma factors, two-component system response regulators, RNA-binding regulators (NusA, NusB, NusG), and other transcription regulators. Of these, 43 regulators were conserved across all three strains as orthologs (**Figure 2D**). To define the conserved regulatory architecture of this core, identify its highest-impact regulators, and reveal the biological functions they coordinate, we next assembled a cross-species RNA-seq compendium spanning all three strains and inferred the core gene regulatory network (GRN) over the conserved regulators.

### 3.3. Integration of cross-species RNA-seq with shared-genome homology yields a cyanobacterial expression compendium for core GRN inference

To identify conserved gene expression architecture and high-impact transcription regulators across the three model cyanobacteria, we assembled a high-quality cross-species RNA-seq compendium spanning PCC 7942, PCC 6803, and PCC 7002. We then used this compendium to infer a core gene regulatory network (GRN) over homologous genes from the shared core genome.

#### A high-quality cross-species expression compendium

We collected raw RNA-seq data from public sequencing repositories including SRA (<u>83</u>), GEO (<u>45</u>) and JGI (<u>84</u>) through 31 January 2024, yielding 1,497 samples across 87 bioprojects (776 samples for PCC 6803, 434 for PCC 7942, and 287 for PCC 7002; **Supplementary Dataset S1)**. After processing the raw data into gene counts (see Methods) and applying a rigorous quality control workflow, we retained species-specific high-quality datasets (Syn7942express, 330 samples; Syn6803express, 548 samples; Syn7002express, 220 samples), which we then merged via shared-genome homology into a single integrated compendium, SynCOREexpress, comprising 1,098 unique expression states across the three species under diverse growth conditions, environmental and genetic perturbations. SynCOREexpress provides a high-quality cross-species expression resource on the shared core genome, enabling direct comparison of conserved gene regulation. Metadata (BioProject numbers, experimental condition, genotype, growth parameters, and quality-control flags) and Log2 TPM counts for all four datasets are provided in **Supplementary Dataset S2.**

#### Core Gene Regulatory Network (GRN) inference and basic network properties

We applied the random forest-based GENIE3 algorithm (<u>65</u>) to the SynCOREexpress compendium, constrained by the 43 conserved candidate regulators identified above, to infer a genome-scale transcription regulatory network. The initial fully connected core GRN covered approximately 96% of the core genome and contained 42 regulators, 1,312 genes, and 55,062 predicted regulatory interactions. To prioritize the highest-confidence TF–gene interactions for downstream analysis, we pruned low-scoring edges to obtain a final trimmed core GRN (**Figure 3A**) of 889 genes (65% of the core genome; **Supplementary Dataset S6-1**), 38 transcription regulators (**Supplementary Dataset S6-2**), and 1,100 TF–gene interactions (**Supplementary Dataset S6-3**). The trimmed network has a median TF degree of 20.5, median TF betweenness of 0.055, average in-degree of 1.24 ± 0.46 (number of TFs acting on a gene), network density of 0.001, and clustering coefficient of 0.08.

**Figure 3.**
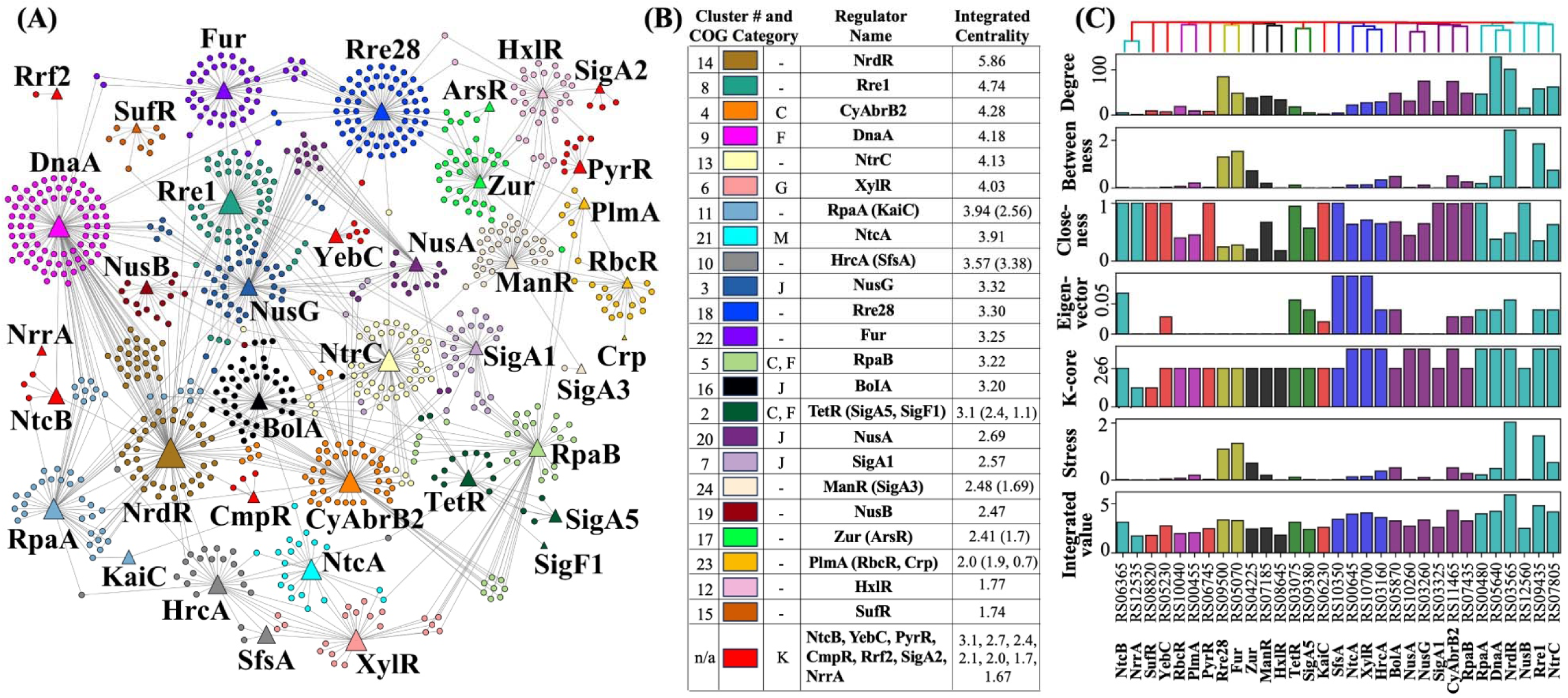
Core GRN of three model cyanobacteria: *S. elongatus* PCC 7942, *Synechocystis* sp. PCC 6803, and *Picosynechococcus* sp. PCC 7002. (A) Interconnected directed graph of the conserved core GRN, comprising 38 transcriptional regulators (triangles, sized proportionally to their Integrated Centrality) and their 851 regulated genes (circles). Nodes are colored by Louvain cluster membership; edges represent predicted TF–gene interactions. **(B)** Louvain cluster identifiers and color scheme, paired with each cluster’s transcriptional regulator(s), enriched COG category, and Integrated Centrality. Parenthetical secondary regulators in the Regulator Name and Integrated Centrality columns indicate additional regulators clustered with the primary regulator. Cluster n/a groups eight regulators with COG category K. **(C)** Six individual centrality measures (degree, betweenness, closeness, eigenvector, k-core, stress) and the Integrated Centrality, calculated for each of the 38 regulators. Regulators along the x-axis are grouped by hierarchical clustering on Jaccard distance reflecting shared regulatory targets. Regulator labels are shown in two rows: locus identifiers (top) and standardized regulator names (bottom). Sigma factor nomenclature is unified across species for visual clarity in cross-species comparison: SigA1 denotes the primary group-1 sigma factor (RpoD1 in PCC 7942 literature), and SigA4–SigA6 denote group-2 alternative sigma factors (RpoD2–RpoD4 in PCC 7942 literature); SigF1 and SigF2 are group-2 sigma factors involved in pilus and biofilm regulation.

With the core GRN established, we next applied topological analysis to identify the conserved regulators with the highest network influence on cyanobacterial gene expression. Although GRN inference from expression data has limited accuracy at the level of individual transcription factor–gene edges, network-level topology reliably recovers biologically meaningful regulatory architecture for cyanobacterial systems (<u>32</u>), and our subsequent cross-validation across three independently inferred species-specific networks provides further independent confirmation of high-influence regulators. As detailed in the following section, this analysis revealed that the highest-influence regulators in the conserved core are predominantly stress-coupled, spanning multi-stress, nutrient, metal, and uncharacterized two-component stress categories.

### 3.4. Topological analysis of the core GRN identifies stress-response regulators as the central elements of the transcription architecture

We analyzed topological properties of the core GRN to identify high-influence conserved regulators, examine the interplay between transcription factors and sigma factors, and characterize the biological functions of their associated gene communities. These analyses establish which regulators occupy the most influential network positions, how stress-responsive regulation is distributed across the conserved core, and which downstream processes these stress-coupled regulators control. Across local, global, and community-aware centrality measures, top conserved regulators of the core GRN are enriched for stress responses including light, redox, heat, osmotic, nutrient, and metal stresses. This observation, quantified below, indicates that stress-response regulators are central elements of the transcription architecture shared among PCC 7942, PCC 6803, and PCC 7002.

#### 3.4.1. Community detection reveals functional specialization, with stress-response and energy-metabolism clusters prominent in the core GRN

We used the Louvain method for community detection, followed by functional enrichment assessment of these communities using categories from the Clusters of Orthologous Genes (COG) database (<u>85</u>) and the KEGG pathway database (<u>72</u>). Of 23 gene clusters (>12 nodes per cluster), 10 were enriched for biological functions (**Figure 3A** and **B**). The enriched communities revealed that the highest-influence regulators in the core GRN partition into two functional groups: stress-coupled regulators controlling major metabolic and biosynthetic processes, and housekeeping regulators controlling translation.

Communities controlled by stress-coupled regulators were enriched for energy and nucleotide metabolism functions. The community associated with CyAbrB2 (an AbrB-family regulator coordinating C/N balance under environmental shifts and dark-to-light transitions (<u>86-88</u>)) was enriched for Energy Production and Conversion (COG category C; *q-value* 2.3×10□□). The communities associated with RpaB (a multi-stress two-component regulator activated by light, redox, and nutrient deprivation (<u>19</u>, <u>89-91</u>)) and TetR together with the co-regulating sigma factors SigA5 and SigF1 were similarly enriched for Energy Production and Conversion (*q-values* 2.5×10□□ and 1.1×10□³, respectively), with secondary enrichment for Nucleotide Transport and Metabolism (COG category F; *q-values* 0.04 and 0.01). The DnaA community, whose regulator couples DNA replication to heat-shock and oxidative-stress responses (<u>66</u>, <u>67</u>, <u>92</u>, <u>93</u>), was likewise enriched for Nucleotide Transport and Metabolism (*q-value* 7.4×10□³). The communities associated with the nitrogen-stress regulator NtcA (activated under N-limitation (<u>94-96</u>)) and the carbohydrate-metabolism regulator XylR (<u>97</u>, <u>98</u>) were enriched for Cell Wall/Envelope/Membrane Biogenesis (COG category M; *q-value* 0.024) and Carbohydrate Transport and Metabolism (COG category G; *q-value* 0.031), respectively.

In contrast, the communities associated with SigA1, NusA, and NusG were enriched for constitutive housekeeping functions: Translation, Ribosomal Structure and Biogenesis (COG category J; *q*-values of 2.6×10□¹¹, 2.5×10□□, and 2.3×10□□, respectively). The community controlled by BolA, a BolA-family regulator that links morphological adaptation and broader physiological responses to multiple stress signals (<u>99-101</u>), was also enriched for Translation, Ribosomal Structure and Biogenesis (COG category J; q-value 7.6×10□³), suggesting that BolA may play a role in coupling stress-driven physiological remodeling to ribosomal and translational adjustments in cyanobacteria.

KEGG enrichment largely agreed with the COG results. The RpaB-regulon community was enriched for Photosynthesis (KEGG map00195; *q-value* 5.9×10□³) and Oxidative Phosphorylation (KEGG map00190; *q-value* 6.9×10□³), consistent with RpaB’s role in coordinating photosynthetic and respiratory gene expression in response to light and redox state (<u>19</u>, <u>89-91</u>).

#### 3.4.2. Multiple network centrality measures consistently identify stress-related transcription regulators among the highest-influence nodes in the core GRN

To identify the conserved regulators with greatest influence on cyanobacterial gene expression, we analyzed local, global, and community-aware network topology measures (**Figure 3C**) for all regulators in the core GRN. In the analyses below, we examine the regulators that emerge at the top of each measure type before turning to the Integrated Centrality measure that combines them.

##### Local Network Topology Measures

Regulators with the highest local centrality (degree, k-core) have the largest number of direct connections to target genes. The transcriptional regulators with the highest degree and k-core values include DnaA (128 direct targets), NrdR (101), Rre28 (84; SYNPCC7942_RS09500), NusG (75), CyAbrB2 (74), NtrC (62), Rre1 (58), BolA (48), Fur (48; SYNPCC7942_RS05070), RpaB (48), and RpaA (46) (**Figure 3B-C** and **Table 2**). Most of these top regulators (DnaA, NrdR, Rre28, CyAbrB2, NtrC, Rre1, BolA, Fur, RpaB, RpaA) are stress-coupled (multi-stress, nutrient, or metal), indicating that stress-responsive regulators are not restricted to small specialized regulons but control broad target sets in the conserved core. The remaining hubs (NusG and the housekeeping regulators discussed above) reflect the constitutive transcription and translation machinery against which this stress-responsive control operates.

**Table 2.**
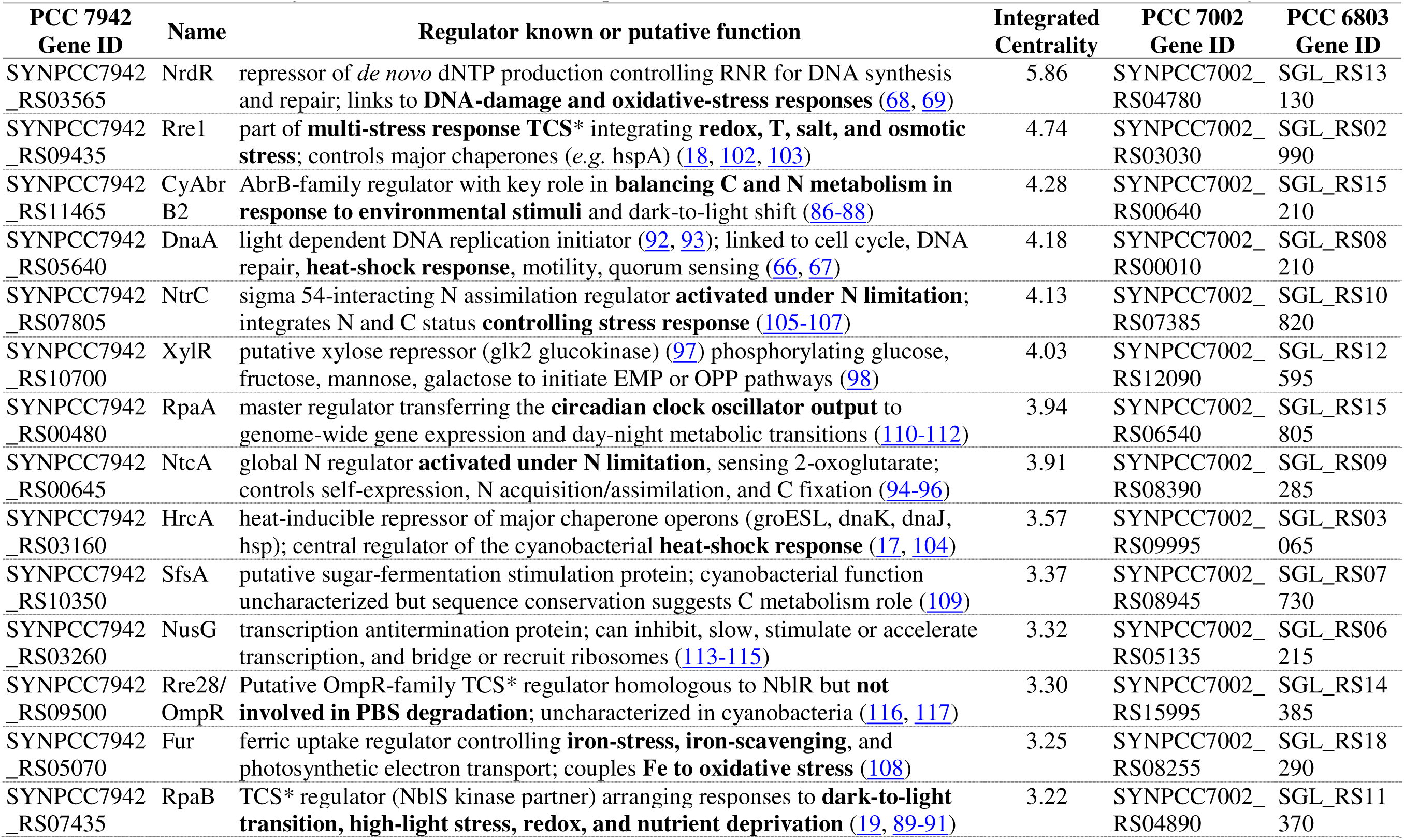
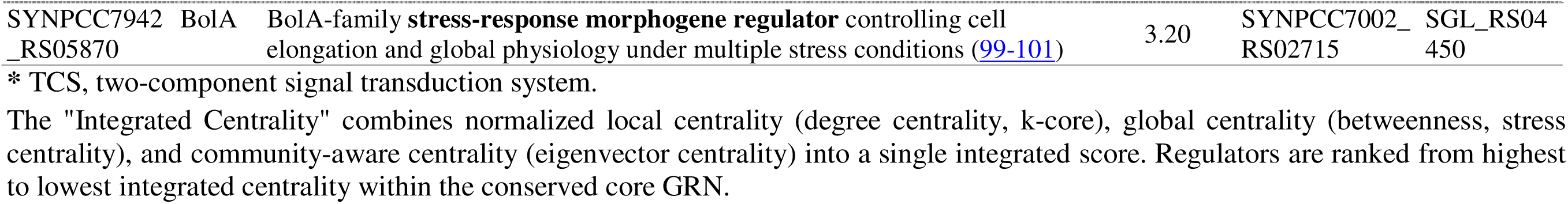
Top high-influence transcriptional regulators in the conserved core GRN, ranked by Integrated Centrality. Regulators ranking among the highest-impact nodes in the conserved core GRN by an Integrated Centrality metric combining local, global, and community-aware network topology measures (Figure 3) are listed for *Synechococcus elongatus* PCC 7942, *Synechocystis* sp. PCC 6803, and *Picosynechococcus* sp. PCC 7002. The 15 regulators shown rank highest in the conserved core GRN by Integrated Centrality; the table provides the current RefSeq locus ID in PCC 7942, the regulator name, a brief functional description with supporting references, and the matched ortholog locus IDs in PCC 7002 and PCC 6803. Of these 15 regulators, 11 (73%) are stress-coupled across multi-stress, nutrient-stress, and metal-homeostasis categories.

##### Global Network Topology Measures

Regulators with the highest global centrality (betweenness, stress, closeness) are positioned to mediate signal flow between functional modules. Top-ranked regulators by these measures include NrdR (betweenness 2.62), Rre1 (2.0), Zur (1.65; SYNPCC7942_RS04225), Rre28 (1.2), NtrC (0.80), CyAbrB2 (0.55), BolA (0.52), DnaA (0.52), HrcA (0.37), RpaB (0.27), and RpaA (0.20). The global ranking features largely the same stress-coupled regulators as local centrality, with two additions: HrcA (heat-shock) and Zur (zinc-stress), neither among the top local hubs. NrdR also moves to the top position by these global measures. Stress-responsive regulators recurring across local and global measures therefore control broad target sets and occupy topological positions for propagating regulatory signals between functional modules of the conserved core.

##### Community-Aware Network Topology Measure

Regulators with the highest eigenvector centrality are connected to other highly connected nodes and are therefore embedded within tightly co-regulated functional modules. The regulators with the highest eigenvector centrality values are XylR (0.095), SfsA (0.095), NtcA (0.095), NtcB (0.067), NrdR (0.056), and TetR (0.056; SYNPCC7942_RS03075). Unlike local and global measures, this ranking is not dominated by broad-spectrum stress regulators; instead, it surfaces regulators of specific functional modules: sugar and carbon metabolism (XylR, SfsA, TetR), nitrogen status sensing (NtcA, NtcB), and DNA-damage and replication stress response (NrdR). The nitrogen and DNA-damage regulators remain stress-coupled, consistent with the local and global rankings, while the sugar and carbon metabolism regulators represent module-embedded controllers tightly integrated into specific functional groups of the conserved core. Together, the local, global, and community-aware measures identify a stress-enriched set of broad-spectrum regulators operating alongside module-embedded metabolic regulators; the Integrated Centrality measure below combines these complementary views.

#### 3.4.3. An Integrated Centrality measure confirms functionally diverse stress-response regulators among the highest-impact nodes in the core GRN

Integrated Centrality combines local, global, and community-aware measures into a single regulator-influence ranking (**Figure 3B** and **3C**). **Table 2** presents the top 15 regulators by this integrated ranking, together with their biological functions, references to characterization studies, and orthologous loci across PCC 7942, PCC 6803, and PCC 7002. Of these 15 regulators, 11 (73%) are linked to stress responses across four categories: multi-stress (light, redox, heat, osmotic), nutrient stress (N, P), metal stress (Fe, Zn), and uncharacterized two-component stress response. The remaining four reflect housekeeping transcription, circadian regulation, and broad metabolic control. This quantitatively confirms the qualitative pattern from the individual measures: stress-response regulators are enriched among the highest-influence positions of the conserved core GRN, with stress-coupling spanning multiple physiological dimensions rather than a single stress type.

Stress-coupled regulators in the core GRN are distributed across four functional categories (**Table 2**). The multi-stress category is the largest, containing six regulators that respond to combinations of light, redox, thermal, and osmotic signals: NrdR (DNA-damage and oxidative stress (<u>68</u>, <u>69</u>)), Rre1 (redox, temperature, salt, and osmotic stress (<u>18</u>, <u>102</u>, <u>103</u>)), HrcA (heat-shock response (<u>17</u>, <u>104</u>)), RpaB (light, redox, and nutrient deprivation (<u>19</u>, <u>89-91</u>)), BolA (multi-stress morphogenic response (<u>99-101</u>)), and DnaA, whose role as a replication initiator is coupled to DNA-damage and heat-shock responses in diverse bacteria (<u>66</u>, <u>67</u>). Three nutrient-stress-related regulators primarily sense nitrogen and coordinate N acquisition with C metabolism: CyAbrB2 (<u>86-88</u>), NtrC (<u>105-107</u>), and NtcA (<u>94-96</u>). CyAbrB2 additionally couples to dark-to-light shifts perturbing C/N balance. Fur is the only metal-stress regulator in the top 15, coupling iron sensing to oxidative stress and photosynthetic electron transport (<u>108</u>). Rre28, an OmpR-family two-component response regulator homologous to NblR but with downstream function in cyanobacteria undefined (<u>18</u>, <u>102</u>, <u>103</u>), represents an uncharacterized stress-response paralog with a substantial regulon.

The four non-stress regulators in the top 15 reflect three additional functional roles. XylR and SfsA are putative regulators of sugar and carbohydrate metabolism (<u>97</u>, <u>98</u>, <u>109</u>). RpaA serves as the master output regulator of the circadian oscillator, transferring clock-driven signals to genome-wide gene expression (<u>110-112</u>). NusG is a transcription antitermination factor with broad transcriptional impact, representing the constitutive transcription machinery against which stress-responsive regulation operates (<u>113-115</u>). Together, the top 15 regulators reveal a conserved core in which stress-response regulators occupy the most influential network positions across multiple stress dimensions, complemented by a smaller set of carbon-metabolism, circadian, and housekeeping regulators.

Across the **centrality measure types** and the integrated ranking, the conserved core GRN emerges as a stress-response architecture: ten of the eleven stress-coupled regulators in the top-15 set (**Table 2**) rank highly by either local or global centrality (or both), and stress-coupling extends across multi-stress, nutrient, metal, and uncharacterized two-component categories rather than concentrating in any single stress type. The conserved transcriptional architecture shared among PCC 7942, PCC 6803, and PCC 7002 therefore concentrates influence in stress-responsive regulators, with circadian, carbon-metabolism, and housekeeping regulators occupying complementary roles. We next examined whether this stress-organized core is paralleled by conserved high-influence regulators across species, and whether species-specific innovations layer additional functional capacity onto it.

### 3.5. Inference of species-specific GRNs and their topological characteristics

Building on the cross-species compendium described above, we inferred species-specific GRNs for PCC 7942, PCC 6803, and PCC 7002 by applying GENIE3 (<u>65</u>) to the corresponding individual datasets Syn7942express, Syn6803express, and Syn7002express (**Supplementary Dataset S2**), and pruning low-weight edges to retain ∼1,100 regulator–gene interactions per network. The basic topological properties of the resulting networks (**Table 3**) include 66, 73, and 81 transcriptional regulators and median regulator degrees of 24.5, 33, and 31 for PCC 7942, PCC 6803, and PCC 7002, respectively. The species-specific networks contain more regulators than the 38 in the conserved core GRN; these additional regulators lack orthologs across all three species and were therefore excluded from the conserved core. We then assessed each network with the same local, global, and community-aware centrality measures applied to the core GRN, allowing direct comparison of high-influence regulators across the conserved core and the three species-specific networks.

**Table 3.** Topological properties of the species-specific gene regulatory networks inferred for *Synechococcus elongatus* PCC 7942, *Synechocystis* sp. PCC 6803, and *Picosynechococcus* sp. PCC 7002. Networks were inferred from species-specific RNA-seq compendia (Syn7942express, Syn6803express, Syn7002express) using GENIE3 with low-weight edge pruning (Methods). Each species-specific GRN includes more regulators than the 38-regulator conserved core GRN because regulators lacking orthologs across all three species are restricted to a single species and were excluded from the conserved core.

|  | PCC 7942 | PCC 6803 | PCC 7002 |
| --- | --- | --- | --- |
| Number of nodes | 897 | 964 | 929 |
| Number of regulators | 66 | 73 | 81 |
| Number of edges | 1,100 | 1,094 | 1,100 |
| Clustering coefficient | 0.096 | 0.034 | 0.048 |
| Median regulator degree | 24.5 | 33 | 31 |
| Median regulator betweenness | 0.0091 | 0.0091 | 0.062 |

### 3.6. Cross-validation across species-specific GRNs highlights stress-response regulators as the most consistently conserved high-influence nodes

To test whether high-influence regulators of the conserved core GRN (Table 2) recur when each species is analyzed independently, we identified top regulators of each species-specific GRN by the same centrality measures and intersected them with the core regulators (**Figure 4**). Across the three species, this cross-validation set comprises 19 regulator instances representing 15 distinct regulators; each species contributes a partially overlapping subset, and not all regulators recur in all three species. Stress-coupled regulators are enriched in this cross-validated set, providing independent support for the stress-organized architecture of the conserved core.

Four regulators recurred as high-influence regulators in two of the three species-specific GRNs (**Figure 4**), the strongest level of cross-validation observed in our analysis. NtcA, the global nitrogen-control regulator activated under N-limitation, ranked among top eigenvector and integrated-centrality regulators in PCC 7942 and PCC 6803, consistent with its established role in coordinating nitrogen acquisition with carbon metabolism (<u>94-96</u>). The sigma 54-interacting nitrogen-assimilation regulator NtrC (SGL_RS10820 in PCC 6803; SYNPCC7002_RS07385 in PCC 7002), which links nitrogen status to broader stress responses (<u>105-107</u>), showed parallel cross-validation in PCC 6803 and PCC 7002. Two multi-stress regulators completed the 2-species set. Rre1 (SGL_RS02990 in PCC 6803; SYNPCC7002_RS03030 in PCC 7002), a multi-stress two-component regulator integrating redox, temperature, salt, and hyperosmotic signals (<u>18</u>, <u>102</u>, <u>103</u>), ranked highly by betweenness and stress centrality in PCC 6803 and PCC 7002; its prominence in these two species, both of which tolerate elevated salinity, is consistent with a role in marine and brackish-niche adaptation, and its appearance among the top integrated-centrality regulators of PCC 7002 further suggests an outsized regulatory role in the species with the broadest known stress tolerance. BolA, a stress-responsive morphogenic regulator that links cell elongation to physiological adaptation under multiple stresses (<u>99-101</u>), ranked among top betweenness and stress-centrality regulators in PCC 7942 and PCC 6803, supporting its proposed role as a global physiological coordinator beyond morphogenesis alone.

Eleven additional regulators ranked highly in only one species-specific GRN while also ranking top in the core GRN. Several of these single-species hits trace patterns of strain-specific stress adaptation. Zur (SYNPCC7002_RS12395), the zinc-uptake regulator, ranked among top community-aware and integrated-centrality regulators of PCC 7002, the only marine cyanobacterium in our analysis, where oceanic zinc is typically limited. ManR (SYNPCC7942_RS07185), the manganese-stress regulator of the ManS-ManR two-component system, was a top local-centrality regulator in PCC 7942, the freshwater species in our set, where manganese homeostasis is shaped by different environmental constraints than in marine systems. Rre28 (SGL_RS14385), an OmpR-family two-component response regulator that has been examined in cyanobacteria but not assigned a defined function (<u>116</u>, <u>118</u>), emerged as a top regulator of PCC 6803 by both betweenness and Integrated Centrality, identifying it as a high-priority candidate for experimental characterization. The remaining single-species top regulators are the multi-stress regulators RpaB and HrcA in PCC 7942 (top by local and global centrality), the carbon-metabolism regulator XylR and the circadian regulator TetR (SYNPCC7942_RS03075) in PCC 7942, the nutrient-stress regulator CyAbrB2 and the circadian master regulator RpaA in PCC 6803, and the multi-stress regulator NrdR and the housekeeping antitermination factor NusG in PCC 7002 (**Figure 4**).

Of the 19 cross-validated ortholog instances (15 distinct regulators) shown in Figure 4, 15 (79%) belong to one of the four stress categories defined in this work: multi-stress, nutrient stress, metal stress, or uncharacterized two-component stress response. Stress-coupled enrichment is consistent across centrality types but most pronounced for global measures (betweenness, stress), where every cross-validated regulator is stress-coupled, and for local centrality (degree, k-core), where stress regulators account for ∼78% of instances. Eigenvector centrality, which captures embedding within tightly co-regulated modules, identifies a smaller, more functionally diverse set: the carbon-metabolism regulator XylR and circadian regulators RpaA and TetR alongside the nutrient-stress regulator NtcA. Several of the stress-coupled cross-validated regulators (notably Rre28, but also Zur, ManR, and several family-level paralogs) remain less well characterized in cyanobacteria than the well-studied core regulators (**Table 2**), identifying them as priority targets for experimental work.

Within each centrality measure, each cyanobacterium has species-specific top regulators overlapping with top regulators of the conserved core GRN (**Figure 4**). Each species also has top regulators by these measures that do not rank highly in the core GRN, reflecting species-specific adaptation and specialization that complement (rather than replicate) the conserved core; these are the focus of the next section.

### 3.7. Species-specific high-influence regulators show broader functional diversity than the conserved core, with stress regulators present but no longer dominant

Beyond regulators that cross-validate between the conserved core and species-specific GRNs (**Figure 4**), each species’ GRN also contains top regulators not ranking highly in the conserved core by the same measure. These species-specific top regulators are shown in **Figure 5** and listed in **Table 4**, organized by species, centrality grouping, and functional category.

**Figure 4.**
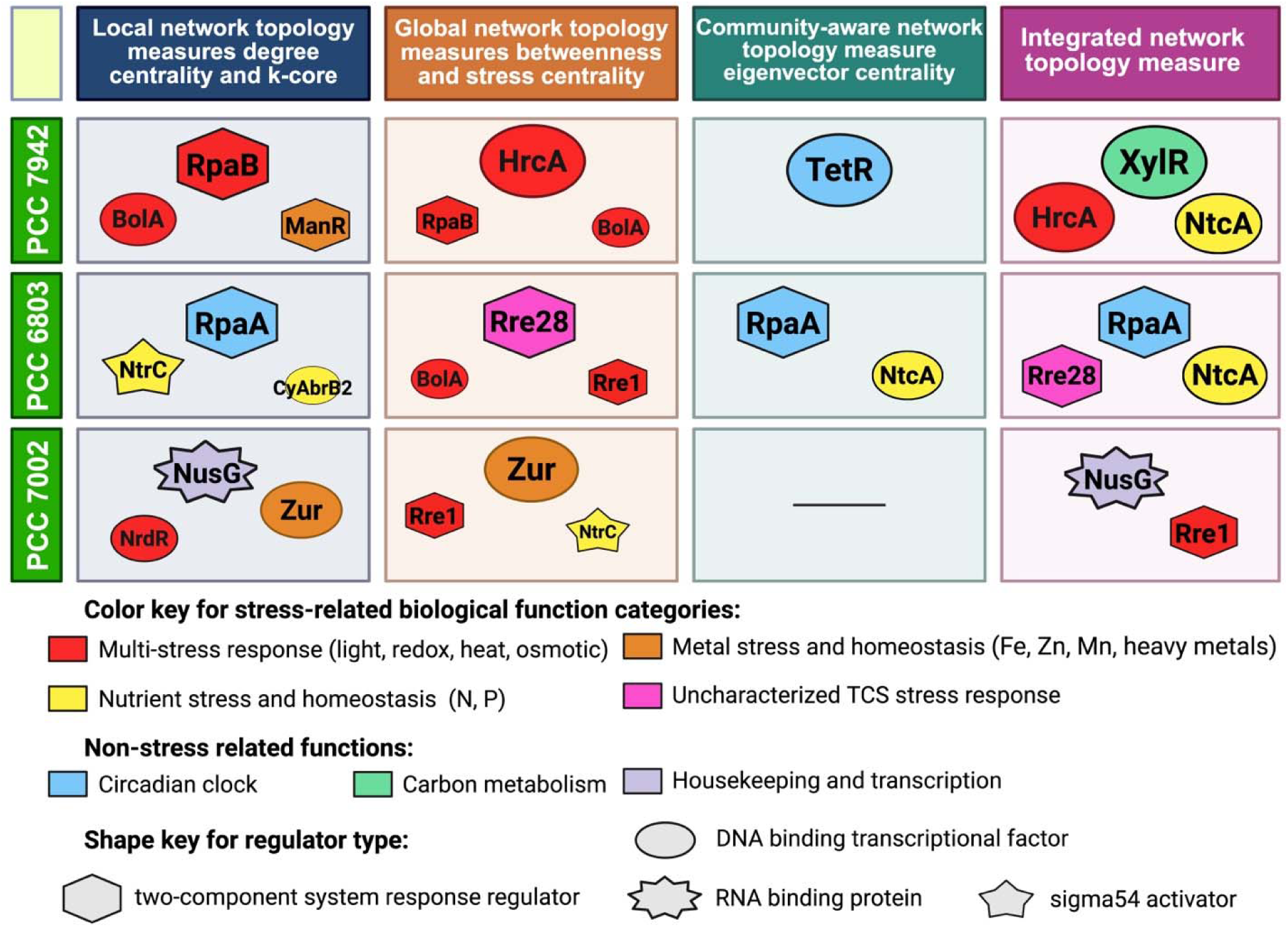
Cross-validation of conserved high-influence regulators shared across the core GRN and species-specific GRNs of *S. elongatus* PCC 7942, *Synechocystis* sp. PCC 6803, and *Picosynechococcus* sp. PCC 7002. Each cell shows regulators ranking highly in both the core GRN and the corresponding species-specific GRN by the indicated centrality measure. Rows correspond to the three species; columns to the four centrality groupings: local (degree, k-core), global (betweenness, stress), community-aware (eigenvector), and integrated. Empty cells indicate no regulator met the cross-validation criterion for that species and measure. Across the three species, the figure shows 19 cross-validated ortholog instances representing 15 distinct regulators; 15 (79%) belong to one of four stress-related functional categories (multi-stress; nutrient stress; metal stress; uncharacterized two-component stress response). Color key: red (multi-stress: light, redox, heat, osmotic), yellow (nutrient stress: N, P), orange (metal stress: Fe, Zn, Mn, heavy metals), pink (uncharacterized two-component stress response), blue (circadian clock), green (carbon metabolism), and purple (housekeeping and transcription). Shape key: ovals, DNA-binding transcription factors; hexagons, two-component system response regulators; star-burst, RNA-binding proteins; stars, sigma-54 activators. Regulator symbols are sized proportionally to the regulator’s normalized centrality score within each cell and ordered top-to-bottom by descending score. Locus identifiers and full functional descriptions are in Table 2. Note: some regulators (ManR, TetR, Zur) appear in cells here by an individual centrality measure but not in **Table 2**, which ranks regulators by Integrated Centrality only; conversely, DnaA, Fur, and SfsA rank top in **Table 2** by Integrated Centrality but not by any individual measure shown here.

**Figure 5.**
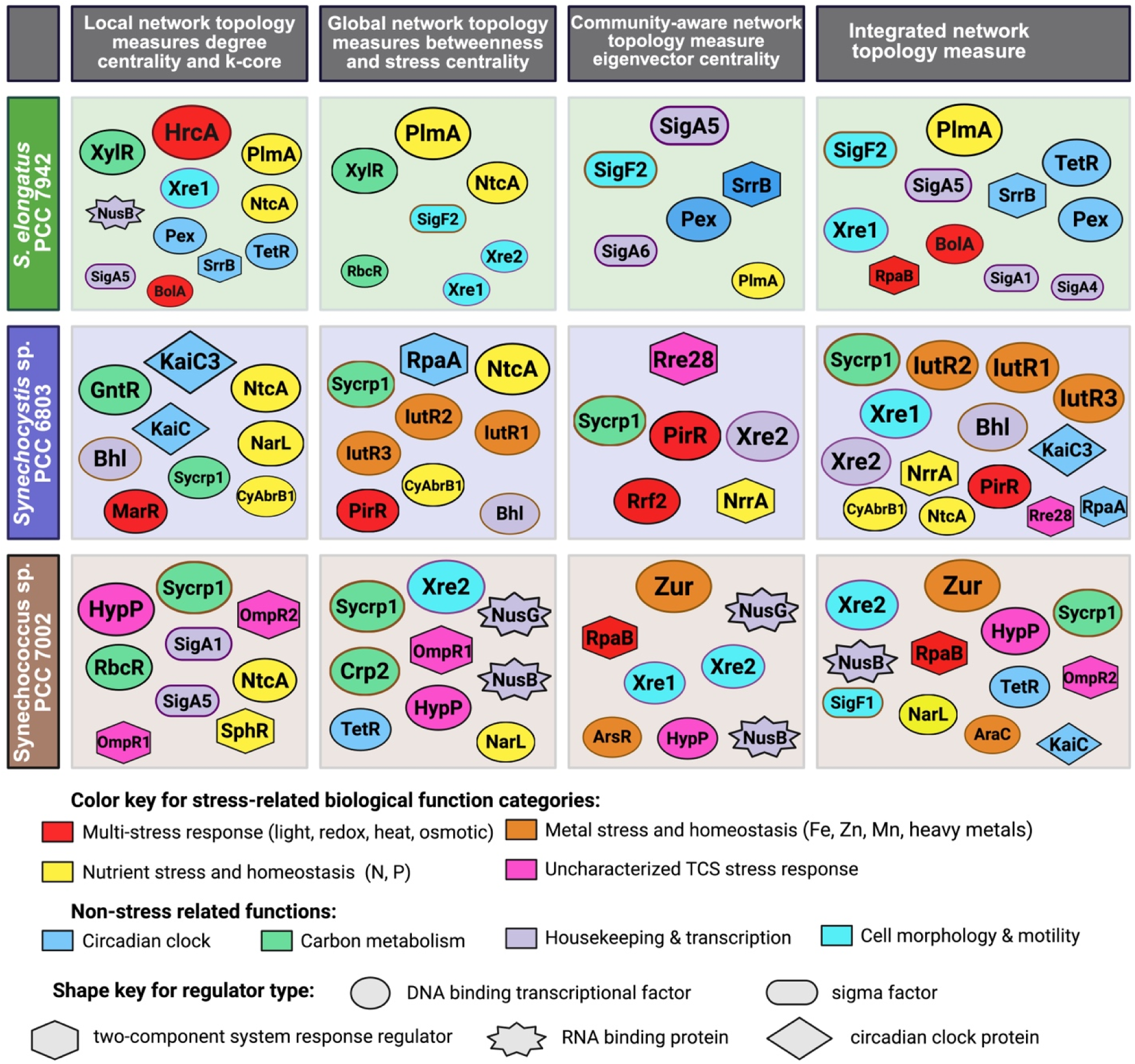
Species-specific high-influence regulators of *S. elongatus* PCC 7942, *Synechocystis* sp. PCC 6803, and *Picosynechococcus* sp. PCC 7002. Each cell shows top regulators of the corresponding species-specific GRN by the indicated centrality measure that did not rank highly in the core GRN by that same measure (the complementary set to Figure 4). Rows correspond to the three species; columns to the four centrality groupings: local, global, community-aware, and integrated. Across the three species, the figure contains 111 cell entries representing 39 distinct regulators, of which 47 (42%) are stress-coupled. Color and shape keys follow Figure 4, with the addition of cyan (cell morphology and motility) and a diamond shape (circadian clock proteins such as the KaiC paralogs). Regulator symbols are sized proportionally to the normalized centrality score within each cell and ordered top-to-bottom by descending score. Naming conventions: three iron-stress AraC-family paralogs in *Synechocystis* sp. PCC 6803 are renamed IutR1, IutR2, and IutR3 following recent published characterization; Xre-family paralogs are labeled Xre1 and Xre2 in PCC 6803 and PCC 7002; OmpR-family paralogs in PCC 7002 are labeled OmpR1 and OmpR2. Sigma factor names are unified across species (SigA1, SigA4–SigA6 for group-1 and group-2 sigma factors; SigF1, SigF2 for pilus/biofilm regulation). Locus identifiers and full functional descriptions are in **Table 4**.

**Table 4.** Top high-influence regulators in species-specific GRNs that do not rank among the top regulators of the conserved core GRN by the same centrality measure. Regulators ranking among the highest-impact nodes in species-specific GRNs (by local, global, community-aware, or Integrated Centrality measures; Figure 5) but not among the top regulators of the conserved core GRN by the same centrality measure (Figure 3, **Table 2**) are listed for *Synechococcus elongatus* PCC 7942, *Synechocystis* sp. PCC 6803, and *Picosynechococcus* sp. PCC 7002. Some regulators are absent from the conserved core GRN entirely; others are present in the conserved core but rank below its top tier by the same centrality measure under which they rank highly in species-specific GRNs. For each regulator, the table provides the current RefSeq locus ID, the regulator name, and a brief functional description with supporting references.

| Locus ID | Name | Regulator known or putative function |
| --- | --- | --- |
| ***Synechococcus elongatus* PCC 7942** |  |  |
| SYNPCC7942_RS00455 | **PlmA** | GntR-family TCS; couples **N-stress response with NtcA-controlled C/N balance** via PII-PipX (119) |
| SYNPCC7942_RS03465 | **Pex** | PadR-family circadian period extender; **tempers kaiA promoter** to fine-tune the oscillator (138, 139) |
| SYNPCC7942_RS02830 | **SrrB \*** | OmpR-family TCS; controlled by RpaA/SasA to follows circadian rhythm (140); putative role in night metabolism (32); homologs in *S. aureus* (redox sensor (141)) and *S. rochei* (antibiotic regulator (142)) |
| SYNPCC7942_RS09740 | **Xre1 \*** | Xre-family DNA-binding protein; uncharacterized; no orthologs in PCC 6803 and PCC 7002 |
| SYNPCC7942_RS09065 | **SigF2** | group-2 $\sigma$ factor; controls **pili and biofilm formation**; 2<sup>nd</sup>-highest Integrated Centrality in PCC 7942 |
| SYNPCC7942_RS03325, RS02895, RS09380 | **SigA1/SigA4 /SigA5** | **large set of $\sigma$ factors among PCC 7942 species-specific top regulators**: primary group-1 SigA1 (RpoD1) and group-2 alternative $\sigma$ factors SigA4, SigA5 (RpoD4, RpoD2 in PCC 7942 literature) |
| ***Synechocystis* sp. PCC 6803** |  |  |
| SGL_RS10605 | **Sycrp1** | cAMP receptor protein; global regulator with targets spanning signal transduction, photosynthesis, CCM, envelope synthesis, DNA repair, phosphate & AA transport (125-128); motility via pili (122, 123) |
| SGL_RS01955, RS01980, RS02020 | **IutR1/IutR2 /IutR3** | AraC-family iron metabolism regulators (formerly PchR/PcrR/PchR); induced by Fe-limitation and H<sub>2</sub>O<sub>2</sub>, suppressed by Fe-excess; putative roles in physiological regulation of Fe homeostasis (129, 130) |
| SGL_RS07535 | **PirR \*** | LysR-family regulator; derepresses pirin ortholog *pirAB* and itself under **salt/osmotic stress** (143) |
| SGL_RS12005 | **CyAbrB1 \*** | AbrB-family paralog of CyAbrB2 with weaker global impact; involved in **nutrient and dark-to-light stress response**; represses C uptake, polysaccharide degradation, and C/N balance (144, 145) |
| SGL_RS00320 | **Xre1 \*** | Xre-family DNA-binding protein on plasmid pSYSM; no orthologs in PCC 7942 and PCC 7002 |
| SGL_RS01625 | **Xre2 \*** | Xre-family DNA-binding protein on plasmid pSYSX; proposed role in antisense RNA regulation |
|  |  | linked to a putative bacterial multiubiquitin system (146). no orthologs in PCC 7942 and PCC 7002 |
| SGL_RS17760 | **KaiC1** | canonical circadian clock protein; **close homolog of KaiC in PCC 7942 and PCC 7002** (131, 132) |
| SGL_RS12430 | **KaiC3** | nonstandard KaiC paralog with the most regulatory targets in PCC 6803; proposed to **regulate day–night transition and dark growth linked to chemoheterotrophic capability** (131, 132) |
| SGL_RS06410 | **Bhl** | HU-family nucleoid protein; tunes **metal stress sensing and tolerance** via DNA topology (147, 148) |
| SGL_RS14385 | **Rre28 \*** | OmpR-family TCS; linked to nutrient and high-light stress via orthology to Synechococcus NblR, but knockout shows normal NblA induction and only delayed bleaching under N starvation (116, 118) |
| ***Picosynechococcus* sp. PCC 7002** |  |  |
| SYNPCC7002_RS12395 | **Zur** | Fur/Zur-family regulator; controls Zn-stress by modulating Zn intake- and homeostasis-related genes under Zn-replete/deplete states (133, 134); broader role likely from high regulatory influence |
| SYNPCC7002_RS08965 | **Sycrp1** | cAMP receptor protein; ortholog of more characterized PCC 6803 Sycrp1 (see above) |
| SYNPCC7002_RS14245 | **HypP \*** | **putative protein on plasmid pAQ7; top regulator in all centrality measures in PCC 7002 GRN** |
| SYNPCC7002_RS11685 | **Xre2** | Xre-family DNA-binding protein; putative homolog of inner-membrane RodZ altering cell shape (135) |
| SYNPCC7002_RS04690 | **OmpR1 \*** | OmpR-family TCS; **putative stress-signal transduction** role, *P. aeruginosa* biofilm/motility (149, 150) |
| SYNPCC7002_RS07760 | **OmpR2 \*** | OmpR-family hybrid TCS sensor kinase/response regulator; **putative role in ion or stress sensing** |
| SYNPCC7002_RS15520 | **NarL \*** | LuxR-family TCS; PCC 6803 KdpE homolog (Sll1592) activated under NH<sub>4</sub><sup>+</sup>/osmotic stress (136); ΔNarL metabolome implicates perturbation of TCA, glycolysis, and pentose phosphate pathway (137) |
| SYNPCC7002_RS15680 | **AraC \*** | AraC-family **on plasmid pAQ1**; putative PchR-like iron/siderophore regulator; distinct from IutR |

**Color key for biological function categories:** red = multi-stress response (light, redox, heat, osmotic); orange = metal stress and homeostasis (Fe, Zn, Mn, heavy metals); yellow = nutrient stress and homeostasis (N, P); pink = uncharacterized two-component system stress response; light blue = circadian clock; green = carbon metabolism; light purple = housekeeping and transcription; cyan = cell morphology and motility.
\* Lack of direct regulator characterization; candidate for experimental work.
The IutR1/IutR2/IutR3 paralogs in PCC 6803 are presented as a single combined row given their close functional similarity; the SigA1, SigA4, and SigA5 sigma factors in PCC 7942 are likewise combined.

Following the centrality groupings of **Figure 4** (local, global, community-aware, integrated), we identified species-specific top regulators of each species’ GRN that do not rank among the top regulators of the conserved core (**Figure 5**). The figure contains 111 cell entries representing 39 distinct regulators, with regulators appearing in multiple cells when ranking highly by more than one measure or when conserved orthologs rank highly in more than one species. Of these 111 entries, 47 (42%) belong to one of the four stress categories: nutrient stress (19 entries), metal stress (10), uncharacterized two-component stress response (10), and multi-stress (8). The remaining 64 entries (58%) span housekeeping transcription, carbon metabolism, circadian regulation, and cell morphology and motility, with the last category appearing for the first time in this analysis. The proportion of stress-coupled regulators in the species-specific top set is therefore substantially lower than in the cross-validated set (79%, **Figure 4**), while non-stress functional diversity is greater. This contrast indicates that the species-specific layer of cyanobacterial transcriptional regulation extends the conserved stress-organized core with broader functional capacity, including cell-shape and motility regulation that is absent from the conserved core.

#### Synechococcus elongatus PCC 7942

The species-specific top regulators of PCC 7942 are led by PlmA, a GntR-family regulator that ranked first by Integrated Centrality and ranked highly by every individual centrality measure. PlmA is proposed to co-regulate nitrogen metabolism with NtcA in response to changing C/N balance via the PII–PipX complex (<u>119</u>), coupling nitrogen-stress response to broader C/N coordination. A second feature of the PCC 7942 species-specific set is an unusually large set of sigma factors among its top regulators, including the group-2 alternative sigma factor SigF2 (which controls pilus and biofilm regulation) and three sigma factors involved in housekeeping transcription (SigA1, SigA4, SigA5; SYNPCC7942_RS03325, RS02895, RS09380) (<u>120</u>, <u>121</u>). Other PCC 7942–specific top regulators include the circadian regulators Pex and SrrB, and the Xre-family DNA-binding protein Xre1, all of which are detailed in **Table 4**.

#### Synechocystis sp. PCC 6803

PCC 6803’s top regulators are led by Sycrp1, the cAMP receptor protein, ranking highest by Integrated Centrality and all individual centrality measures. Initially identified as a regulator of motility and pili formation (<u>122</u>, <u>123</u>), Sycrp1 is now recognized as a global regulator analogous to its *E. coli* homolog CRP, which controls 574 targets (<u>124</u>). Its computationally predicted and partially validated regulatory targets in PCC 6803 include the carbon-concentrating mechanism, photosynthesis, signal transduction, cell envelope synthesis, DNA repair, and phosphate transport (<u>125-128</u>). This study supports a global regulatory role for Sycrp1.

PCC 6803 is also distinguished by three iron-regulated AraC-family paralogs (among top four nodes) that we rename here as IutR1, IutR2, and IutR3 (formerly PchR), all induced by iron limitation and H O exposure, and suppressed by iron excess (<u>129</u>, <u>130</u>). A third notable feature is KaiC3, a nonstandard paralog of the canonical circadian clock protein KaiC that has the highest number of regulatory targets in the PCC 6803 network and is proposed to regulate growth in darkness, consistent with PCC 6803’s chemoheterotrophic capability (<u>131</u>, <u>132</u>). Other PCC 6803–specific top regulators including PirR, CyAbrB1, KaiC1, two Xre-family paralogs (Xre1 and Xre2), and the HU-family histone-like DNA-binding nucleoid-associated protein Bhl, which modulates heavy-metal (Ni, Co, Zn, Hg) sensing and tolerance (details shown in **Table 4).**

#### Picosynechococcus sp. PCC 7002

Two PCC 7002 species-specific top regulators stand out. Zur, a Fur-family zinc-uptake regulator (<u>133</u>, <u>134</u>), ranked highest by community-aware and Integrated Centrality measures, consistent with the importance of zinc homeostasis in this marine cyanobacterium where seawater zinc is typically depleted. The high network influence of Zur in PCC 7002 suggests regulatory roles beyond canonical zinc transport. HypP, a hypothetical protein encoded on plasmid pAQ7 (SYNPCC7002_RS14245), ranked highly by all four centrality measures but has no published functional characterization, making it a high-priority experimental candidate. A third PCC 7002–specific regulator, Xre2 (SYNPCC7002_RS11685), is a putative functional homolog of the inner-membrane protein RodZ involved in cell-shape control (<u>135</u>), representing the cell-morphology category that distinguishes the species-specific layer from the conserved core. Other PCC 7002–specific top regulators including Sycrp1, two uncharacterized OmpR-family regulators (OmpR1, OmpR2), the ammonium- and osmotic-stress responsive NarL impacting central carbon metabolism (<u>136</u>, <u>137</u>), and a plasmid-encoded AraC are detailed in **Table 4**. The species-specific regulators identified across the three cyanobacteria suggest concrete targets for cross-species regulatory engineering, which we explore below.

### 3.8. From network topology to transcription machinery engineering: candidate regulators for cross-species biotechnology

The conserved core and species-specific regulatory layers identified above translate into candidate engineering targets for cyanobacterial biotechnology. Sigma factor engineering and transcription factor modification have enhanced complex phenotypes in bacteria, yeast, and microalgae (**Table 5**), but their application in cyanobacteria has been limited by the absence of a systematic regulator catalog with network positions. Most high-influence regulators in our analysis are stress-coupled and therefore map naturally onto the central engineering challenges: tolerating abiotic stresses of outdoor and industrial cultivation, redirecting carbon from storage and biomass toward target products, and coordinating production with native temporal regulation. Below, we organize candidate regulators around these three application areas.

**Table 5.** Representative examples of transcription machinery engineering in cyanobacteria, microalgae, and other microbial systems. Endogenous overexpression and knockdown studies in cyanobacteria are listed first, followed by heterologous expression (cross-species transfer) within cyanobacteria, and cross-kingdom precedents from microalgae, bacteria, and yeast as proof of principle for global transcription engineering

| Organism | Regulator target<br>(gene) (type) | Mod. | Outcome |
| --- | --- | --- | --- |
| **Endogenous engineering in cyanobacteria** |  |  |  |
| *Synechocystis* sp.<br>PCC 6803 | SigE ($\sigma 70$ grp 2) | OE / KD | OE: 50–120% ↑ glycogen/sugar catabolism (*glgP*, *glgX*, *opcA*, *zwf*, *gnd*, *talC*); 250% ↑ PHB; ↑ H<sub>2</sub> (micro-oxic); ↓ respiration and photosynthesis; aberrant cell division. KD: ↓ sugar catabolism; ↓ *hox* and *nifJ* fermentative genes (micro-oxic) (153-157) |
| *Anabaena* (*Nostoc*)<br>sp. PCC 7120 | SigE ($\sigma 70$ grp 2) | OE / KD | OE: ↑ heterocyst frequency, ↑ nitrogenase. KD: ↓ glycolysis and PPP; ↓ heterocyst-specific genes; delayed differentiation (172, 173) |
| PCC 7120 | SigC ($\sigma 70$ grp 2) | KD | Delayed heterocyst development; ↓ OPPP and *ntcA* (174) |
| PCC 7120 | OrrA (TCS RR) | OE / KD | OE: ↑ sucrose synthesis genes; $\sim 2$-fold ↑ sucrose. KD: ↓ sucrose synthesis; loss of post-dehydration re-growth (158, 159) |
| PCC 7120 | NrrA / Rre37 (OmpR) | OE | Enhanced heterocyst formation (175, 176) |
| PCC 7120 | NtcA (global) | KD | Loss of sucrose induction under N deprivation (95) |
| PCC 7120 | FurC / PerR (Fur-family) | OE | Activation of heterocyst development, bicarbonate uptake, glycogen phosphorylase, N fixation, OPPP-driven reducing power, PS machinery maintenance (163) |
| PCC 7120 | CyAbrB1 / *calA* (AbrB-family) | OE / KD | OE: heterocyst formation abolished under N depletion. KD: ↑ heterocyst formation under nitrate via ↑ *hetP* and *hepA* (177) |
| *S. elongatus* PCC 7942 | SigF1 ($\sigma 70$ grp 2) | OE | Biofilm formation in otherwise planktonic culture; ↑ matrix gene expression and secretion; pilus assembly blocked but cells remain transformable (151, 152) |
| PCC 7942 | SigF2 ($\sigma 70$ grp 2) | OE | Pili and planktonic growth unaffected; biofilm not inhibited; non-transformable (152) |
| *Synechocystis* sp.<br>PCC 6803 | CyAbrB2 (*sll0822*) (AbrB-family / NAP) | KD | Deregulated C/N metabolism; growth inhibition; $\sim 100\%$ ↑ free fatty acids; ↑ glycogen; *hox* and *nifJ* less repressed (aerobic) and more repressed (micro-oxic) (88, 157, 167, 178) |
| PCC 6803 | LexA (AbrB-like) | KD | ↓ *hox* hydrogenase activity; de-repressed FA synthesis ( ↑ FAs); de-repressed *ndhF3*, *cmpA*, *sbtA* at high CO<sub>2</sub>; impaired CO<sub>2</sub> -limitation response (11 C-related genes affected) (168, 179-182) |
| PCC 6803 | PrqR (*slr0895*)<br>(TetR/AcrR) | KD | ↑ growth rate and ↓ ROS under high light via de-repression of *sodB* and *prqA*; activates glucose metabolism through *sigE* and *gap2* (162) |
| *Nostoc punctiforme* | SigC, SigJ (σ70 grp 2) | KD | Disrupted hormogonium development; loss of cell-size reduction, heterocyst loss (183) |
| Cyanobacteria PCC 6803, PCC 7002, PCC 7942, PCC 7120 | NrrA (OmpR RR) | OE / KD | OE (PCC 6803, N-replete): activated glycogen → PHB, TCA, ornithine flux; ↑ glycogen catabolism, glycolysis, PHA synthesis; OPPP repressed. KD across strains: ↓ glycogen catabolism, glycolysis, OPPP, respiration; ↑ glycogen accumulation; delayed heterocysts; ↓ N-limitation survival (164, 165, 173, 176, 184-186) |
| **Heterologous expression in cyanobacteria** |  |  |  |
| *Anabaena* sp. PCC 7120 | SigJ from *Nostoc* HK-01 (σ70 grp 3) | HE | Acquired desiccation tolerance; 3.2-fold ↑ EPS release (160, 161) |
| **Cross-kingdom proof of principle** |  |  |  |
| *Escherichia coli* | RpoD (Pri σ70) | gTME | Enhanced ethanol tolerance via global transcriptome reprogramming (3) |
| *S. cerevisiae* | SPT15 (TATA) | gTME | Enhanced ethanol tolerance; altered global gene expression (4) |
| *C. glutamicum* | SigA / SigB | OE / KD | 2-fold (SigA OE) and 2.5-fold (SigB KD) ↑ carotenoid (decaprenoxanthin) (187) |
| *Escherichia coli* | IrrE (from *D. radiodurans*) | HE | Enhanced tolerance to ethanol, salt, and oxidative stress (188) |
| *Nannochloropsis* spp.; *Chlorella* sp. | bZIP TFs | OE | 50–120% ↑ FAME content; 16.2-fold ↑ extracellular lipid (NobZIP1, ChIP-confirmed); ↑ lipogenic genes (11, 189, 190) |
| *N. salina* | NsbHLH2 | OE | 49% ↑ FAME productivity under N-limited continuous culture (191, 192) |
| *N. gaditana* | ZnCys TF (Zinc-finger) | KO | 2-fold ↑ lipid productivity via redirected C from protein to lipid (12) |
| *N. salina* | AtWRI1 (*A. thaliana*) | HE | 64% ↑ FAME yield; ↑ *PPDK*, *LPL*, *LPGAT1*, *PDH* (193) |
|  | (AP2-type) |  |  |
| *C. reinhardtii* / *C. ellipsoidea* | GmDof4 (*G. max*)<br>(Dof-type) | HE | 2-fold ↑ total lipids in *C. reinhardtii*; 53% ↑ lipid content in *C. ellipsoidea* (*ACCase* ↑) (194, 195) |
| *C. reinhardtii* | CrDof (Dof-type) | OE | 23.24% ↑ total fatty acid content (196) |

**Abbreviations:** OE, overexpression; KD, knockdown; KO, knockout; HE, heterologous expression; gTME, global transcription machinery engineering; TCS RR, two-component system response regulator; NAP, nucleoid-associated protein; PPP, pentose phosphate pathway; OPPP, oxidative pentose phosphate pathway; PHB, polyhydroxybutyrate; PHA, polyhydroxyalkanoate; FAME, fatty acid methyl ester; FA, fatty acid; ROS, reactive oxygen species; EPS, extracellular polymeric substance.

Several precedents demonstrate that transcription machinery engineering is technically feasible in cyanobacteria, producing phenotypes spanning morphology, stress tolerance, and central metabolism (**Table 5**). Overexpression of the group-2 sigma factor SigF1 in *S. elongatus* PCC 7942 converted the planktonic strain into biofilm-forming culture by activating matrix-gene expression and blocking pilus assembly (<u>151</u>, <u>152</u>), showing a single sigma-factor modification can switch a multicellular phenotype. The paralog SigF2 produced a distinct outcome, leaving pili and planktonic growth intact while rendering cells non-transformable (Suban et al., 2024), showing that even closely related sigma factors can be tuned to different traits. In *Synechocystis* sp. PCC 6803, overexpression of the group-2 sigma factor SigE upregulated sugar catabolism and oxidative pentose phosphate pathway genes 1.5–2.2-fold, raised intracellular acetyl-CoA, and increased polyhydroxybutyrate production roughly 2.5-fold (<u>153-157</u>), a precedent we revisit below as a carbon-partitioning lever. In *Anabaena* sp. PCC 7120, manipulating the two-component response regulator OrrA increased sucrose synthesis approximately two-fold under normal conditions and conferred post-dehydration regrowth ability, while its loss abolished both phenotypes (<u>158</u>, <u>159</u>), demonstrating that a single regulator can simultaneously control a stress-tolerance trait and a bioproduction-relevant metabolic output. Cross-species transfer between cyanobacteria has also been demonstrated: heterologous expression of the group-3 sigma factor SigJ from the desiccation-tolerant *Nostoc* HK-01 into PCC 7120 conferred desiccation tolerance and a 3.2-fold increase in extracellular polysaccharide release (<u>160</u>, <u>161</u>). These precedents establish that single-regulator interventions can deliver specific, complex phenotypic outcomes in cyanobacterial hosts.

#### Engineering stress tolerance for industrial cultivation

Outdoor and large-scale cyanobacterial cultivation is limited by combined high-light, oxidative, thermal, and nutrient stresses that depress photosynthetic efficiency and culture stability. The multi-stress regulators central to the conserved core, including Rre1, RpaB, HrcA, and Rre28 **(Table 2**, **Figure 4)**, are natural candidates for re-tuning these responses across strains. Direct engineering precedent already exists for the high-light axis: knockdown of the TetR/AcrR-family regulator PrqR in *Synechocystis* sp. PCC 6803 raised growth rate and lowered reactive oxygen species under high light through de-repression of *sodB* and *prqA*, while simultaneously activating glucose metabolism via *sigE* and *gap2* (<u>162</u>). A second cyanobacterial precedent comes from the Fur-family peroxide-response regulator FurC (PerR) in *Anabaena* sp. PCC 7120, whose overexpression coordinately activated heterocyst development, bicarbonate uptake, nitrogen fixation, oxidative pentose phosphate pathway flux, and photosystem maintenance (<u>163</u>), illustrating that a single broad regulator can bundle stress-protective and productivity-relevant phenotypes. The strong centrality of Zur in *Picosynechococcus* sp. PCC 7002 (**Figure 5**) is a complementary species-specific entry point: as a marine strain adapted to zinc-poor seawater, PCC 7002 may carry Zur-coordinated regulatory adaptations transferable to freshwater hosts for seawater-based cultivation. These examples define a tractable design space for stress-tolerance engineering: network position nominates the targets; existing precedents demonstrate the route.

#### Engineering carbon partitioning

Redirecting fixed carbon from storage and biomass toward target products is a central goal of cyanobacterial biotechnology; several regulators governing this partitioning are already validated engineering levers. The best-developed precedent is SigE in *Synechocystis* sp. PCC 6803: overexpression raises acetyl-CoA pools and increases polyhydroxybutyrate production ∼2.5-fold (<u>153-155</u>), and NrrA acts on a parallel pathway converging on the same flux toward PHB, the tricarboxylic acid cycle, and the ornithine cycle (<u>164</u>, <u>165</u>). Because SigE has not been identified in *S. elongatus* PCC 7942 (<u>166</u>), heterologous expression of PCC 6803 SigE in PCC 7942 is a concrete cross-strain transfer proposal combining our species-specific finding with an established route. CyAbrB2, a top-tier regulator in the conserved core (**Table 2**, **Figure 4**), provides an additional carbon-partitioning lever: its inactivation in PCC 6803 derepresses fatty-acid biosynthesis, roughly doubles free fatty-acid accumulation, and coordinates hydrogenase expression during fermentation (<u>157</u>, <u>167</u>). Knockdown of LexA in PCC 6803 has been similarly used to derepress fatty-acid biosynthesis and activate the bidirectional hydrogenase (<u>168</u>), broadening the carbon-partitioning toolkit to include hydrogen as a co-product. Finally, the central position of NtcA and NtrC in the conserved core (**Table 2**, **Figure 4**) points to a high-payoff opportunity: tuning these nitrogen-stress regulators to uncouple product induction from growth arrest, a trade-off bounding productivity in nutrient-stress-induced bioproduction, recently broken by single-regulator knockdown in *Nannochloropsis gaditana* (<u>12</u>).

#### Engineering temporal control of bioproduction

Cyanobacteria possess one of biology’s best-characterized circadian systems, and recent work in *S. elongatus* PCC 7942 shows the subjective-night phase carries an intrinsically more productive metabolic state, with carbon flux redirected from storage and cell division toward target products such as sucrose (<u>169</u>). This natural separation defines a second engineering opportunity: rather than constitutively driving production at the cost of growth, regulators governing the circadian output state can in principle be tuned to align induction with the favorable window or extend it. Our analysis identifies RpaA as the strongest candidate: the master circadian output regulator in cyanobacteria, cross-validated as a top regulator of the conserved core **(Table 2**, **Figure 4**), sitting at the interface between the Kai oscillator and downstream metabolic programs. KaiC, the central clock component, occupies the same regulatory cluster (**Table 2**) and provides a complementary handle on the oscillator itself; the PCC 6803-specific paralog KaiC3 (**Figure 5**) is a species-specific transfer candidate that may broaden the temporal repertoire of recipient strains. Auxiliary circadian regulators recovered as species-specific high-influence nodes in PCC 7942, particularly PlmA (proposed to co-regulate C/N balance with NtcA via the PII–PipX complex), Pex, and SrrB (**Figure 5**), define a wider toolkit for fine-tuning circadian metabolic output. Coordinating heterologous pathways with these native regulators offers a route to growth–production–harvest cycles in which the most costly steps are temporally separated rather than competing for shared carbon and reducing-power pools.

Several considerations shape how these candidates translate from network position to experimental phenotype. Successful cross-species transfer depends on conservation of target promoters and transcription factor binding sites in the recipient strain, absence of regulatory conflicts with native circuits, and topological compatibility between donor and recipient networks. The strongest cases for predictable transfer are therefore the conserved-core regulators of **Table 2** and **Figure 4**, whose orthologs and binding-site syntax are present in all three strains; species-specific candidates from **Figure 5** carry higher uncertainty but also the potential for larger phenotypic gains where the recipient lacks the underlying regulatory layer. Two priorities emerge directly from our analysis. First, the highest-centrality conserved-core regulators that lack engineering precedent, particularly Rre1 and Rre28, should be characterized experimentally to establish their downstream phenotypes before deployment as engineering targets. Second, extending the analysis to additional cyanobacterial species, including *Anabaena* sp. PCC 7120 and ecologically distinct genera such as *Prochlorococcus*, *Nostoc*, and *Acaryochloris*, would broaden both the conserved-core inventory and the catalog of species-specific innovations available for transfer. Together with the precedents of **Table 5**, the regulators identified here define a tractable, stress-anchored design space for cyanobacterial transcription engineering under environmental stress.

Beyond the regulator-level findings, this work yields a set of reusable resources for cyanobacterial systems biology. SynCOREexpress, the cross-species expression compendium of 1,098 quality-controlled expression states curated from 87 bioprojects across three species (**Supplementary Dataset S2**), the multi-pipeline curated regulator inventories for each strain (**Supplementary Dataset S5**), the conserved core GRN over the 1,362-gene tri-homologous core (**Supplementary Dataset S6**), and the three species-specific GRNs are released for reuse, refinement, and integration into downstream analyses. The Integrated Centrality framework introduced here is independent of cyanobacterial biology and generalizes to other microbial systems where GRN inference from public expression data is tractable.

Several considerations bound the present analysis. First, the conserved core and species-specific GRNs are inferred from expression-level associations using GENIE3, not from experimentally validated regulator-target binding events; while network-level topology recovers biologically meaningful regulatory architecture despite limited per-edge accuracy (<u>32</u>), individual edges should not be interpreted as established regulatory relationships. Second, GENIE3 or alternative computational methods identifies regulators correlated with target expression but does not by itself establish causal direction or transcription-factor binding-site occupancy, so directionality of individual interactions remains to be experimentally tested. Third, the high-influence sets analyzed here use top-tier cutoffs that are heuristic by nature; while the cross-species consistency of the top tier (**Figure 4**) supports the underlying ranking, the precise membership of the highest-impact set depends on the chosen threshold. Fourth, three strains capture only part of cyanobacterial regulatory diversity, and the analysis does not include nitrogen-fixing filamentous lineages (*Anabaena*, *Nostoc*), oligotrophic ocean specialists (*Prochlorococcus*), or far-red-pigmented lineages (*Acaryochloris*); extending the framework to these would broaden both the conserved core inventory and the catalog of species-specific innovations. The regulators and ranks reported here are therefore best understood as comparative computational predictions that prioritize experimental targets; validation by (<u>170</u>), DAP-seq (<u>171</u>), regulator knockout or overexpression, and targeted promoter-binding assays remains essential before engineering deployment.

## 4. Conclusions

This study confirmed a two-layer hypothesis of cyanobacterial regulatory architecture, with stress-response regulators enriched among the highest-influence positions of the conserved core and among the network positions most consistently influential across species; species-specific regulators broadened this picture into carbon-metabolism, circadian, morphology, and housekeeping functions. The Integrated Centrality score introduced here proved effective at identifying regulators of joint topological influence and offers a generalizable framework for ranking regulators in other microbial systems. As multi-omics datasets for cyanobacteria continue to grow, machine-learning approaches combining multi-omics integration with network topology offer a natural next step for resolving regulator function at scale, refining engineering candidates, and enabling the engineered cyanobacterial strains that sustainable bioproduction will require.

## ACKNOWLEDGMENTS

This research was supported by the Predictive Phenomics Initiative Laboratory Directed Research and Development Program at Pacific Northwest National Laboratory (PNNL), and U.S. Department of Energy, Office of Science program Biopreparedness Research Virtual Environment (BRaVE) Initiative award to PNNL (81832). Rudolph DiMura, Robert Li, and David Anderson were supported in part by the U.S. Department of Energy, Office of Science, Office of Workforce Development for Teachers and Scientists (WDTS) under the Science Undergraduate Laboratory Internships Program (SULI). PNNL is operated by Battelle for the DOE under Contract DE-AC05-76RL01830.

## Data availability statement

Data is provided within the manuscript or Supplementary Dataset files.

## Authors agreement to authorship and submission

All persons designated as author agree to submit the manuscript for peer review.

## Author contributions

P.B. and M.C. – funding acquisition and project administration; P.B. conception and design; D.A. and P.B. – data collection and pre-processing; R.D., R.L., D.A., Z.J. and P.B. formal analysis of data; R.D., R.L. and P.B – visualization; R.D. and P.B. – writing original draft; P.B. and M.C. – writing review and editing.

## Declaration of Generative AI and AI-Assisted Technologies in the Writing Process

During the preparation of this work, the authors used Anthropic’s Claude to assist with editing and refining manuscript prose, ensuring terminology consistency, verifying that supplementary data files correspond to in-text references, and enhancing overall readability. The authors subsequently reviewed and edited all content and accept full responsibility for the accuracy and integrity of the publication.

## Statement of informed consent, human/animal rights

No conflicts, informed consent, human or animal rights applicable.

